# Unbalanced Perturbation Dynamics For Cell Fate Design

**DOI:** 10.64898/2026.06.30.735555

**Authors:** Qiangwei Peng, Yuchuan Wang, Jianzhe Li, Xinyu Wang, Yao Xiao, Peijie Zhou

**Author notes:** These authors contributed equally to this work (Qiangwei Peng, Yuchuan Wang, Jianzhe Li, Xinyu Wang, Yao Xiao).

## Abstract

Large-scale single-cell perturbation sequencing provides an unprecedented opportunity to construct virtual cells for the in silico simulation of cellular responses and the inverse design of optimal interventions. However, most perturbation-response models treat cellular responses primarily as mass-preserving shifts in transcriptomic state, whereas single-cell perturbation measurements are inherently unbalanced: the recovered endpoint population is shaped by technical sampling as well as biological perturbation-induced proliferation, apoptosis and selection. Here we introduce U-Pert, an unbalanced generative framework that learns condition- and context-dependent perturbation dynamics from unpaired single-cell snapshots. U-Pert jointly models transcriptomic state transitions and cell-number dynamics, enabling scalable and robust forward prediction of unseen perturbations and contexts, as well as inverse design to screen for desired genetic or pharmacological interventions that achieve user-defined transcriptomic or population-level outcomes. Across controlled simulations, genetic perturbation benchmarks, sciPlex3 drug responses and PBMC cytokine perturbations, U-Pert predicts unseen responses, captures both molecular and abundance changes, and performs inverse design for target gene-expression programs and cell-type compositions. These results show that cell abundance is an integral component of the perturbation phenotype, providing a mass-aware framework for virtual-cell modeling and perturbation cell fate design.

## Introduction

Cell fate decisions arise from coordinated changes in gene expression, differentiation, proliferation and cell death [1–3]. A central goal of modern cell biology is therefore not only to observe these changes, but to predict how a cell population will respond when a gene, pathway or drug is perturbed. The emerging concept of an *in silico* virtual cell aims to support this goal by using computational models to prioritize experiments and anticipate cellular responses across biological contexts [4]. For such models to be biologically useful, however, perturbation responses cannot be simply reduced to transcriptomic state changes among recovered cells. Perturbations can indeed alter the abundance and composition of the endpoint population through effects on proliferation, apoptosis and selection [5]. Ignoring this abundance axis can confound perturbation analysis by forcing expansion, depletion or selective survival to be explained only as movement in transcriptomic state space.

Single-cell perturbation technologies, including Perturb-seq and sciPlex, now connect targeted genetic or chemical interventions with transcriptome-wide readouts at single-cell resolution [6, 7]. These assays have revealed gene functions, regulatory programs and genotype–phenotype relationships across rich perturbation landscapes [8–11]. In addition, the resulting datasets also expose the unbalanced nature of perturbation responses: different conditions can yield markedly different numbers and compositions of recovered cells, reflecting both technical factors such as sampling, capture efficiency and experimental design [12] and biological effects of the perturbation itself. At the same time, the space of possible perturbations, doses, combinations, exposure times and biological contexts remains far larger than can be exhaustively tested. Because most single-cell assays are destructive, the cells measured before and after perturbation are not the same cells. This creates a need for models that can infer how an unperturbed population gives rise to a perturbed outcome from unpaired snapshots, while allowing both transcriptomic states and population mass to change.

Substantial progress has been made in modeling single-cell perturbation responses at the gene expression level. For instance, representation-learning approaches learn latent spaces for counterfactual state prediction across cells, conditions and drug perturbations [13–17]. Graph- and regulatory-network-based models use biological structure to predict or simulate genetic interventions [18–20]. Optimal-transport, causal and flow-based approaches infer distribution-level response mappings from unpaired single-cell data [21–24]. Recent foundation, language-informed and perturbation-specific models have further expanded the scale and conditioning capacity of response prediction [25–36].

Despite this breadth, most current methods still treat perturbation response primarily as a change in gene-expression state and leave population mass fixed or implicit. This mismatch is particularly consequential when perturbations induce cell killing, expansion or selection, or when balanced transport forces distributional differences into implausible cross-cell-type matching. Moreover, existing models are largely formulated for forward prediction, whereas many biological and therapeutic questions require inverse design, i.e. identifying genetic or pharmacological interventions that steer cells toward user-defined outcomes, from target differential-expression programs to desired cell-type compositions or abundance phenotypes. Recent benchmark studies have also shown that perturbation prediction remains sensitive to dataset structure, confounding factors and may underperform the simple baselines [5, 37, 38], demonstrating the need for models that better match the biological and experimental structure of perturbation data.

Here we introduce U-Pert (Unbalanced Perturbation), an unbalanced generative framework for learning perturbation-induced cell-state and cell-abundance dynamics from unpaired single-cell data. U-Pert learns condition- and context-dependent perturbation dynamics that couple transcriptomic state transitions with cellular abundance change. This formulation allows the model to predict responses to unseen perturbations and biological contexts while also supporting inverse design to screen for desired genetic or pharmacological interventions that achieve user-defined transcriptomic or population-level outcomes. The unbalanced modeling is particularly important for population-level inverse design, because targets such as cell-type proportions or counts depend on perturbation-induced expansion and depletion rather than transcriptomic state transitions alone. For perturbations specified by discrete identities, such as target genes or drug compounds, U-Pert performs design by optimizing a continuous perturbation representation and retrieving real candidates by embedding similarity; for continuous variables such as dose, it can search the perturbation space directly. Across controlled simulations with known ground truth and broad genetic, pharmacological and cytokine perturbation datasets, U-Pert predicts unseen responses, captures both molecular and abundance changes, and performs inverse design for target gene-expression programs and cell-type compositions. By treating perturbation responses as unbalanced changes in both cellular state and population structure, U-Pert provides a framework for perturbation biology in which virtual cells can move beyond response prediction toward the rational design of cell-fate outcomes.

## Results

### Overview of U-Pert

External biological perturbations, such as gene knockouts [6, 9] or drug treatments [7], leave an immediate transcriptomic signature in single cells, most visibly through the up- or down-regulation of differentially expressed genes (DEGs) (Fig. 1a). This transcriptomic readout has been the primary target of many perturbation-response models [13, 15, 18, 21–24, 29, 31–35]. Across the datasets analyzed, however, perturbations also produced substantial differences in the number of recovered cells (Fig. 1b Left). Because many of these experiments began with controlled initial cell inputs, the resulting endpoint differences reflect an abundance signal beyond initial loading, although capture efficiency, sampling and screening design can also contribute. Perturbations are indeed known to alter proliferative capacity through cell-cycle regulation or apoptosis induction [5, 39, 40] (Fig. 1b Right). These observations motivate modeling perturbation responses as coupled endpoints, combining transcriptomic state changes with condition-dependent variation in recovered cell abundance (Fig. 1b).

**Fig. 1.**
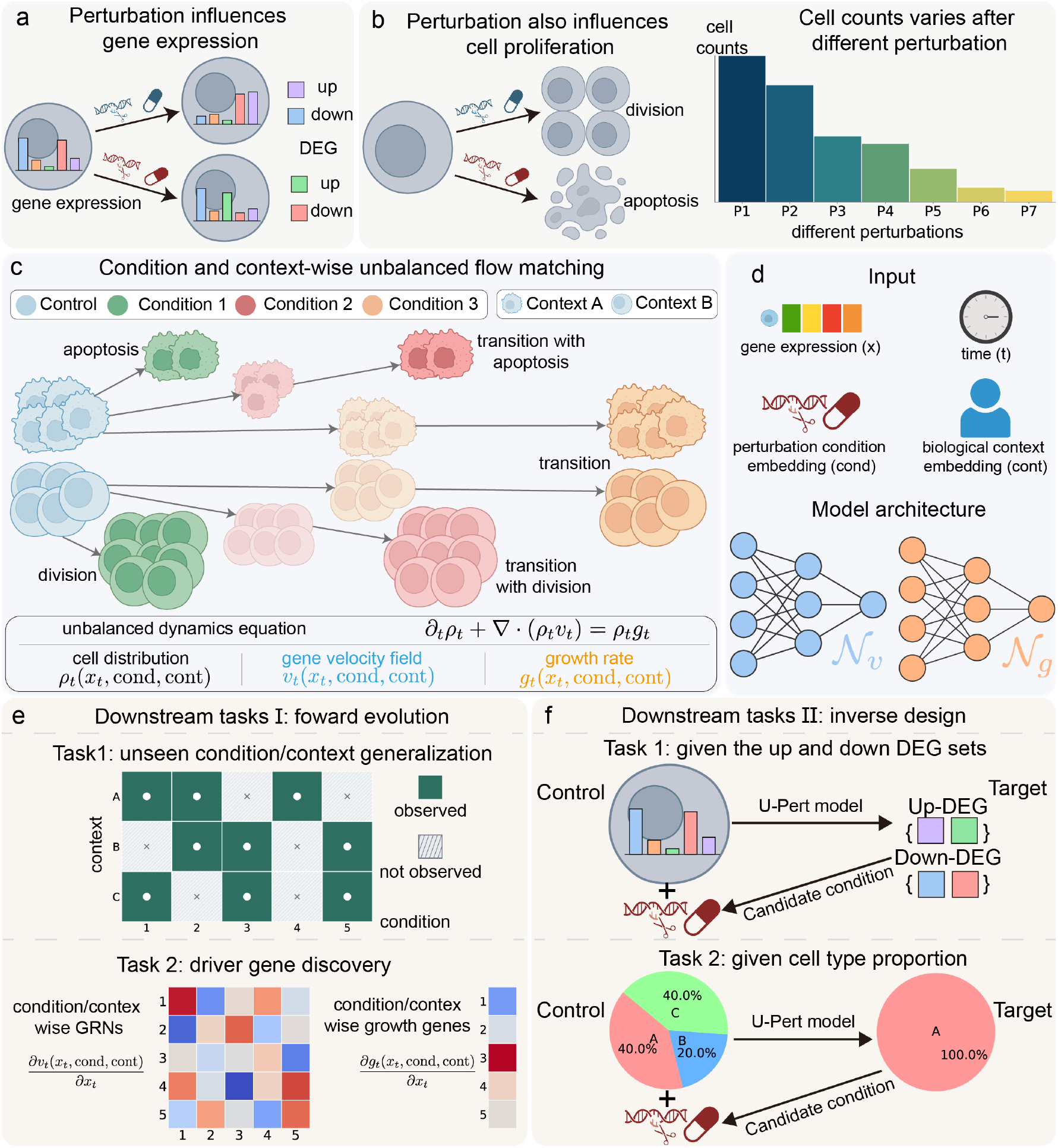
Overview of the U-Pert framework a. External perturbations (e.g., drug treatments or genetic modifications) induce transcriptomic shifts, driving the up- or down-regulation of DEGs. **b**. Different perturbations lead to substantial variations in total cell counts (left), driven by altered cellular proliferation and apoptosis capacities (right), highlighting the necessity of jointly modeling expression and population size changes. **c**. Schematic of the condition- and context-wise unbalanced flow matching. U-Pert seamlessly maps initial control distributions to various perturbed states across distinct biological contexts. The continuous state transition is governed by an unbalanced dynamics equation (transport-with-growth PDE), where the gene velocity field (*v*_*t*_) and growth rate (*g*_*t*_) account for transcriptomic trajectories and varying cell counts (division and apoptosis), respectively. **d**. Model inputs and neural network architecture. The framework takes gene expression (*x*_*t*_), time (*t*), and learned embeddings for condition (cond) and context (cont) to parameterize the velocity (*N*_*v*_) and growth (*N*_*g*_) networks. **e**. Forward evolution downstream tasks. Top: U-Pert generalizes to unseen conditions or contexts to predict unobserved cellular responses. Bottom: Calculation of the Jacobian of the velocity field and the gradient of the growth rate enables the mechanistic discovery of condition-specific GRNs and growth-driver genes. **f**. Inverse design downstream tasks. Top: Given a target transcriptomic state defined by a specific DEG set, U-Pert traces backward to prioritize the candidate perturbation condition required to achieve it. Bottom: By explicitly modeling cell birth-death dynamics, U-Pert can prioritize the candidate perturbation condition needed to orchestrate a target macroscopic cell type proportion or count.

To capture such coupled processes, we develop U-Pert as a condition- and context-wise unbalanced flow matching framework. U-Pert models the temporal evolution of cell populations by coupling transcriptomic state transitions with changes in cell abundance (Fig. 1c and Methods). The state of a cell population is represented as a time-dependent distribution over gene-expression profiles, conditioned on both the perturbation condition and the biological context. Here, the perturbation **condition** can represent a specific gene knockout, drug treatment or dose, whereas the biological **context** can represent covariates such as cell line, tissue type or donor (Fig. 1d; Methods).

Unlike standard balanced optimal transport, which preserves total mass, U-Pert uses an unbalanced transport-with-growth formulation to allow cell populations to expand or shrink during the response. In this formulation, the gene velocity field describes how cells transition through transcriptomic state space, whereas the growth rate function describes how cellular mass changes along the response. Biologically, for a given condition and context, the gene velocity field captures perturbation-induced gene-expression trajectories, while the growth rate function explicitly models changes in cell abundance caused by division and apoptosis (Fig. 1c,d).

A major challenge in modeling single-cell perturbation responses is the destructive nature of sequencing, which yields highly unpaired unbalanced data. Consequently, one cannot observe the same cell both before and after perturbation. U-Pert addresses this challenge by learning response dynamics from unpaired control and perturbed snapshots. For each perturbation condition, it constructs an unbalanced matching between the control and perturbed cell distributions, and then learns condition- and context-dependent velocity field and growth rate from these matched population-level responses (Methods). This allows U-Pert to infer both transcriptomic state transitions and abundance changes directly from static unbalanced single-cell perturbation measurements.

After U-Pert captures the unified perturbation dynamics across diverse conditions and contexts, it enables a suite of downstream tasks categorized into forward prediction and inverse design (Fig. 1e,f and Methods).

#### Forward Prediction

Based on the learned velocity fields and growth rates, U-Pert provides predictive and interpretative capabilities (Fig. 1e).

1. Unseen condition/context generalization. By learning representations for both perturbation conditions and biological contexts, U-Pert can generalize to settings that were not directly observed during training. It simulates forward responses to predict cell distributions and transcriptional states for zero-shot scenarios, including responses to unseen perturbations and biological, such as a held-out gene/drug, or a held-out cell line absent from the training data (Fig. 1e Top and Methods).
2. Driver gene discovery. U-Pert also allows the learned dynamics to be interrogated mechanistically. The gene-expression velocity field can be used to infer condition/context-specific gene regulatory relationships that drive state transitions. In parallel, the growth rate function can be used to identify genes associated with cellular proliferation or apoptosis, linking molecular features to population-level abundance changes (Fig. 1e Bottom and Methods).

#### Inverse Design

A feature of U-Pert is its ability to perform inverse design, reversing the perturbation-to-phenotype workflow to identify interventions for desired biological outcomes (Fig. 1f).

1. Targeting transcriptomic states. Given a desired outcome defined by a set of DEGs, U-Pert can trace the dynamics backward to search the embedding space for the optimal perturbation condition, such as a specific drug or genetic modification, that is predicted to drive control cells toward the target transcriptional state (Fig. 1f Top and Methods).
2. Targeting cell type proportions and counts. By modeling perturbation responses as unbalanced dynamics, U-Pert can use macroscopic population outcomes as inverse-design objectives. It therefore designs perturbations not only toward desired transcriptional states, but also toward target cell-type proportions or absolute cell counts by allowing population mass to expand or contract through the growth field. Given a target composition, such as enriching a rare therapeutic cell population or eliminating a malignant cluster, U-Pert infers the external perturbation condition predicted to achieve this goal (Fig. 1f Bottom and Methods).

### U-Pert enables forward prediction and inverse design of perturbation responses in a controlled dynamical system

We first evaluated U-Pert to recover both components in a controlled synthetic system, where the true perturbation dynamics were explicitly defined (Fig. 2a). The system consists of three genes (*G*_1_, *G*_2_, *G*_3_), whose expression levels (mRNA *X*_1_, *X*_2_, *X*_3_) are governed by an underlying gene regulatory network (GRN) coupled with apoptosis dynamics. Specifically, *G*_1_ and *G*_2_ form a toggle-switch circuit characterized by mutual inhibition, with both genes also exhibiting positive autoregulation. The third gene, *G*_3_, acts as an upstream regulator that suppresses both *G*_1_ and *G*_2_ while being activated by *G*_2_. In addition, *G*_3_ directly promotes apoptosis, thereby linking gene expression dynamics to cell survival dynamics. The transcription rates of *X*_1_ and *X*_2_ are parameterized by *ρ*_1_ and *ρ*_2_, respectively, while the apoptosis rate is controlled by *α* (Methods). In our experiments, only these three dynamical parameters are perturbed, with all other parameters fixed throughout.

**Fig. 2.**
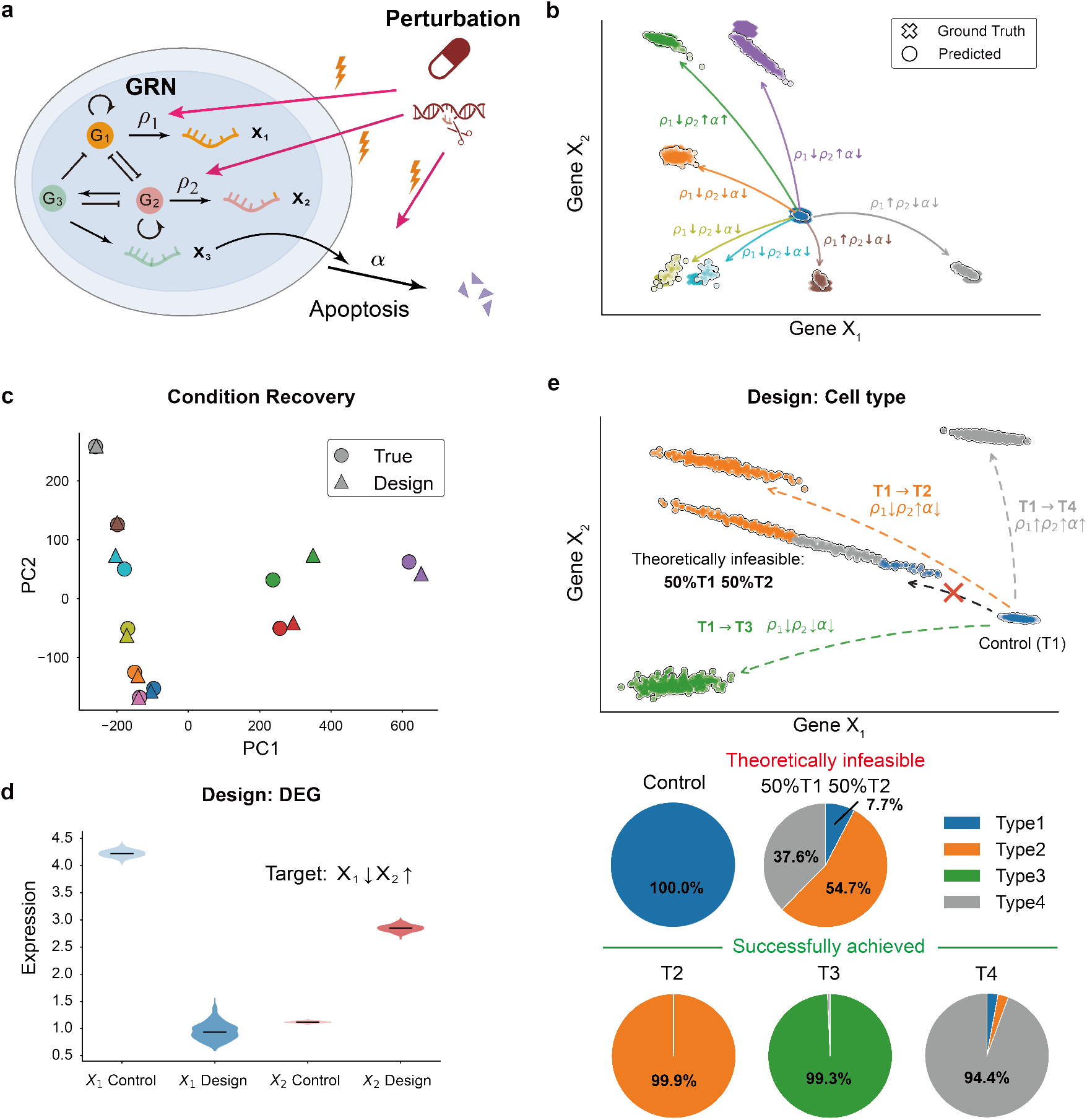
U-Pert enables forward prediction and inverse design of perturbation responses in a controlled dynamical system. a. Schematic of a gene regulatory network (GRN) governing gene expression (*X*_1_, *X*_2_, *X*_3_) and apoptosis dynamics. Perturbations act on transcription rates (*ρ*_1_, *ρ*_2_) and the apoptosis rate (*α*), while other dynamical parameters (not shown) are held constant. **b**. Forward prediction of virtual cells under perturbations in gene expression space (*X*_1_– *X*_2_). Colors denote cell populations, with blue indicating the control and other colors representing perturbed states. Arrows indicate transitions from the control, with parameter changes encoding perturbation effects. Predicted populations closely match ground-truth distributions. **c**. Condition recovery, where designed conditions accurately recapitulate the true perturbation conditions, visualized via a comparison in the PCA1–PCA2 space obtained from the projection of three kinetic parameters: transcription rates (*ρ*_1_, *ρ*_2_) and the apoptosis rate (*α*). **d**. Inverse design of differential gene expression, generating cell populations that satisfy target constraints (*X*_1_ ↓, *X*_2_ ↑). **e**. Inverse design of cell-type composition, demonstrating controllable transitions and proportions; feasible targets are achieved, whereas theoretically infeasible designs are correctly identified.

To evaluate forward prediction under unseen perturbations, we defined a control population of 1,163 cells under a fixed parameter setting and generated perturbed populations by varying (*ρ*_1_, *ρ*_2_, *α*) (Fig. 2b). Evaluation on 100 unseen perturbation conditions shows that U-Pert-predicted populations closely match the ground-truth perturbed populations after projection onto the (*X*_1_, *X*_2_) expression space. Representative examples from several selected perturbation conditions are shown in Fig. 2b, illustrating that U-Pert faithfully captures cellular responses to perturbations.

Beyond forward prediction, U-Pert further enables recovery of underlying perturbation conditions via inverse design. Using target perturbed populations with ground-truth parameters, we inferred the corresponding perturbation conditions from the cellular states. Across 100 unseen perturbation conditions, the inferred conditions achieved an average cosine distance of 0.09 from the true conditions. After projection into PCA space, paired inferred and true perturbation parameters showed strong overlap, indicating accurate recovery of the underlying dynamical parameters (Fig. 2c). For visualization clarity, we show 10 representative perturbation conditions selected from the full evaluation set in this PCA projection.

To further design the desired cell fates, we considered two inverse-design tasks enabled by U-Pert. At the gene level, U-Pert infers perturbations that induce prescribed differential expression patterns, such as decreasing *X*_1_ while increasing *X*_2_, resulting in clear shifts in expression distributions relative to the control (Fig. 2d). At the cell-type composition level, U-Pert enables control over cell-type proportions, where cell types are defined by dominant gene expression patterns (Methods). Starting from a control population dominated by T1 cells, U-Pert successfully generates populations enriched for T2, T3, or T4 cells, achieving final proportions of 99.9%, 99.3%, and 94.4%, respectively (Fig. 2e).

Of note, this controllability indeed did not come from arbitrarily forcing prescribed target compositions. U-Pert respected the intrinsic dynamical constraints of the underlying gene regulatory system. For instance, a 50% T1 and 50% T2 composition cannot be robustly achieved by a single perturbation applied to the fixed initial population. Mechanistically, this behavior is consistent with the toggle-switch structure of the GRN, in which T1 and T2 correspond to two competing stable attractors driven by mutually inhibitory *X*_1_ and *X*_2_ programs. Perturbations attempting to balance the two fates inevitably place part of the population near the separatrix and the associated saddle-like transition region between the two basins of attraction. Cells in this weakly committed regime do not fully converge to either stable fate and are therefore classified as the intermediate state T4. Consequently, U-Pert remains faithful to the attractor geometry of the underlying dynamical system, revealing biologically constrained design limits instead of generating physically implausible population mixtures.

Overall, rather than merely matching endpoint clouds, U-Pert can recover both imposed state dynamics and imposed survival constraints in the synthetic perturbation datasets.

### U-Pert improves transcriptomic and cell-abundance prediction across genetic perturbation benchmarks

We next evaluated the predictive benefit of the unbalanced formulation in real biological datasets. Large-scale single-cell genetic perturbation screens provide a stringent benchmark because perturbation responses must be predicted for unseen genetic interventions, while the recovered endpoint populations vary substantially across conditions. In pooled Perturb-seq experiments, cells are assigned genetic perturbations and then profiled after perturbation-specific growth, death and selection have acted on the population [6, 8, 9, 11]. Consistent with this unbalanced experimental setting, widely-used benchmarking perturbation datasets such as Norman [11], Adamson [8] and Replogle K562 [9] all showed substantial variation in recovered cell abundance across perturbation conditions (Fig. 3a). Because pooled screens profile recovered cells rather than a perfectly controlled output population for every perturbation, these count differences should be interpreted cautiously: some variation may reflect capture, sampling or experimental design as well as biology. Nevertheless, the abundance structure showed biological organization, as high-abundance Norman conditions were enriched for apoptosis regulation, cell proliferation, protein stabilization, hematopoiesis and oxygen transport programs (Fig. 3b). These observations define an observed perturbation response that is intrinsically unbalanced, and U-Pert is naturally suited to this setting because it explicitly models both transcriptomic transport and condition-dependent cell-abundance change [41, 42].

**Fig. 3.**
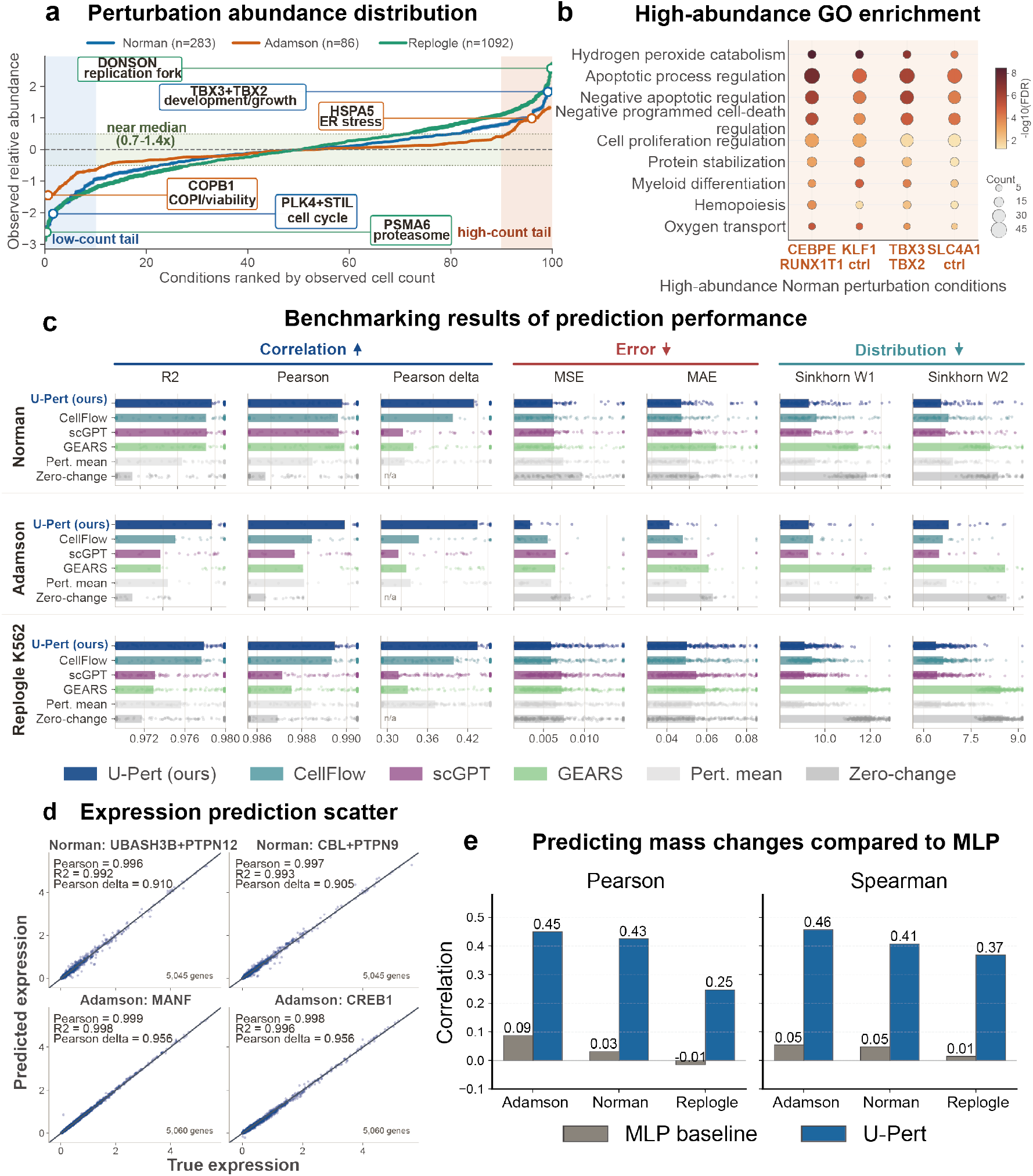
Benchmarking U-Pert on genetic perturbation datasets. **a**. Norman, Adamson and Replogle K562 datasets exhibit substantial imbalance across perturbation conditions. Conditions are ranked within each dataset by observed cell count, and abundance is shown relative to the dataset median. **b**. Gene Ontology enrichment of significantly changed genes in high-abundance Norman perturbation conditions, showing significant enrichment of growth- and cell-death-related biological programs. **c**. Benchmarking of U-Pert against CellFlow, scGPT, GEARS, perturbation-mean and zero-change baselines across Norman, Adamson and Replogle K562. Bars show the mean across held-out perturbation conditions, and points show individual condition-level values. Correlation metrics include *R*^2^, Pearson correlation and Pearson delta, where Pearson delta evaluates correlation of expression changes relative to control. Error metrics include MSE and MAE. Distributional metrics are *W*_1_ and *W*_2_ distances. **d**. Representative gene-level prediction examples for held-out Norman and Adamson perturbation conditions. Each point represents one gene, with true mean expression on the x-axis and U-Pert-predicted mean expression on the y-axis. **e**. Prediction of perturbation-associated mass changes across all conditions. Pearson and Spearman correlations between observed and predicted mass ratios are shown for Adamson, Norman and Replogle, comparing U-Pert with an MLP baseline.

To determine whether an abundance-aware formulation improves genetic-perturbation prediction, we benchmarked U-Pert on unseen perturbation conditions from Norman, Adamson and Replogle K562 using standard perturbation-response metrics (Fig. 3c; Methods). The comparison included representative perturbation-prediction baselines from different modeling families: the flow-matching model CellFlow [23], the single-cell foundation model scGPT [26], the graph-based perturbation predictor GEARS [18], as well as perturbation-mean and zero-change empirical baselines. Across complementary expression metrics, U-Pert showed consistent improvements or competitive performance across all three datasets. The clearest gain was observed for perturbation-induced expression changes, where mean Pearson delta reached 0.636 in Norman, 0.865 in Adamson and 0.434 in Replogle, compared with 0.618, 0.661 and 0.399 for CellFlow. U-Pert also maintained strong absolute-expression and distributional accuracy, indicating that the unbalanced formulation did not compromise conventional gene-expression prediction. Representative unseen conditions further showed calibrated gene-level predictions in both Norman and Adamson, with predicted mean expression closely matching the observed perturbed means and with strong agreement in expression changes relative to control (Fig. 3d). U-Pert therefore retained robust transcriptomic prediction performance while introducing an additional mass readout.

To evaluate this abundance component directly, we compared the predicted mass ratio after perturbation with the observed target-to-source abundance ratio for each condition and benchmarked this readout against a supervised MLP mass baseline using the same condition information. U-Pert consistently improved mass-ratio correlation over the MLP baseline across Adamson, Norman and Replogle (Fig. 3e). Pearson and Spearman correlations increased from 0.088 and 0.054 to 0.450 and 0.458 in Adamson, from 0.031 and 0.047 to 0.425 and 0.407 in Norman and from −0.015 and 0.014 to 0.247 and 0.368 in Replogle K562. This result suggests that the learned growth field provides a useful condition-level abundance readout that is not captured by balanced transcriptomic predictors alone.

Taken together, these benchmarks clarify the value of unbalanced modeling in pooled genetic screens. Although recovered cell number reflects both biological and technical sources of variation, modeling condition-dependent mass helped U-Pert better match the measured perturbation endpoint, improving transcriptomic prediction beyond merely balanced state-only predictors.

### U-Pert captures drug-induced state changes and cell-abundance changes in sciPlex3

Genetic perturbation benchmarks provide evidence that U-Pert improves prediction across unseen interventions. To examine a setting in which abundance change is a more explicit component of the perturbation phenotype, we next investigated chemical perturbations, where dose and cell-line identity can produce transcriptional state changes, cell survival effects or both. We used sciPlex3 [7] to evaluate U-Pert’s ability to jointly recover transcriptional state and abundance changes in real drug responses. The dataset included 188 compounds, three cancer cell lines, four dose levels and two time points (Fig. 4a). Each held-out condition was defined by a tuple of drug, cell line, dose and time. Predictions were generated from matched control cells.

**Fig. 4.**
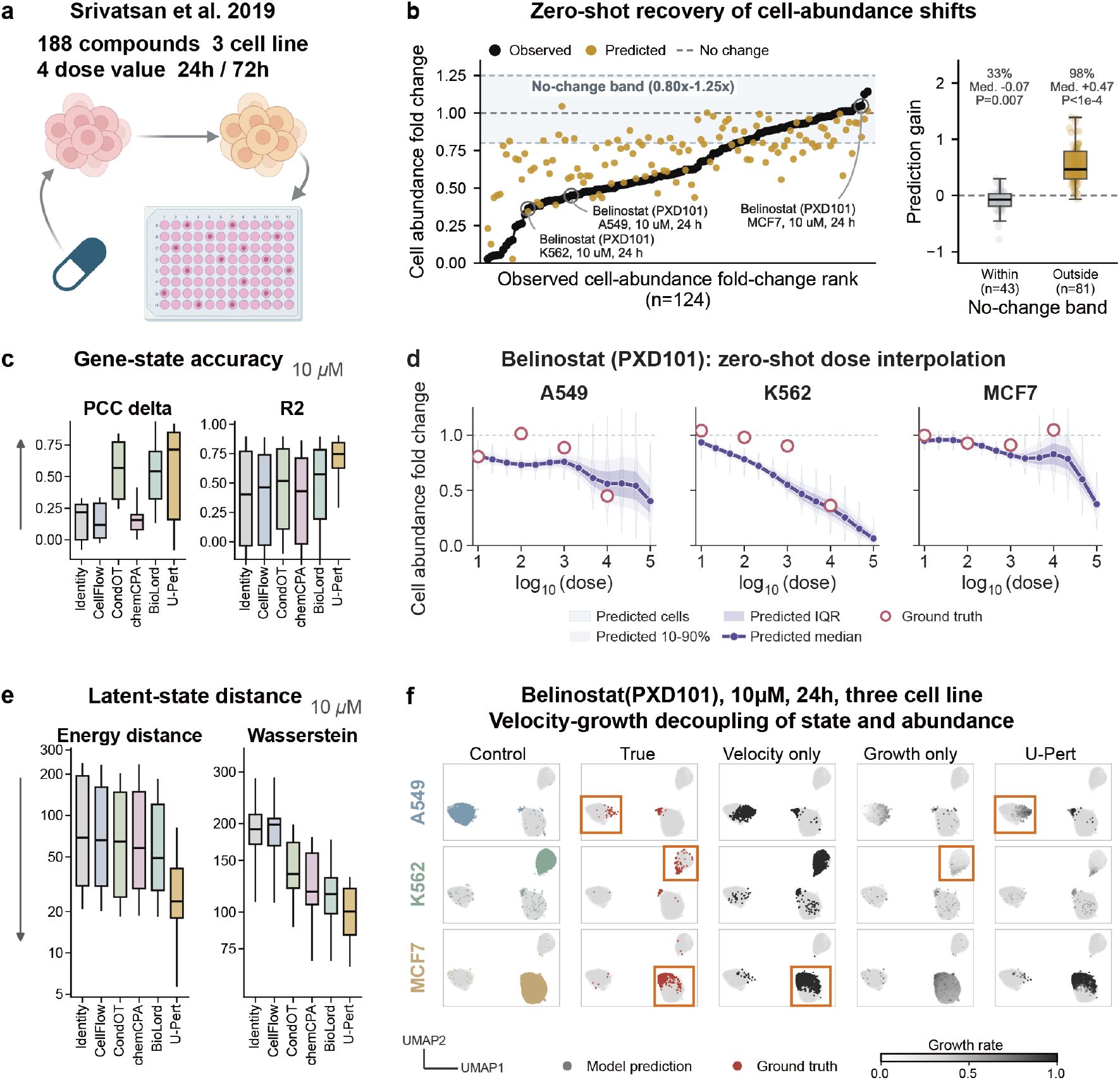
U-Pert jointly predicts transcriptional response and cell-abundance change in sciPlex3. **a**. Overview of the sciPlex3 dataset. It contains 188 compounds profiled across A549, K562 and MCF7 cells, four dose levels and 24 h or 72 h time points. **b**. Recovery of cell-abundance shifts across 124 held-out drug, cell-line, dose and time tuples. The left panel shows observed and predicted cell-abundance fold changes after ranking conditions by observed fold change. The right panel shows prediction gain over a no-change baseline, computed as the reduction in absolute log2 fold-change error. Conditions are grouped by whether the observed fold change lies within the no-change band from 0.8 to 1.25. **c**. Gene-state benchmark across the same held-out tuples. Boxplots compare Identity, CellFlow, CondOT, chemCPA, BioLord and U-Pert using delta-expression correlation and gene-level R2. Higher values indicate better agreement with the observed response. **d**. Zero-shot dose interpolation for belinostat (PXD101) at 24 h in A549, K562 and MCF7 cells. Predicted particle-level mass distributions are shown across a log-spaced dose grid, with observed abundance fold-change points overlaid. **e**. Latent-distribution benchmark across the same held-out tuples. Boxplots show latent energy distance and latent Wasserstein distance, for which lower values indicate better agreement with the observed perturbed distribution. **f**. Velocity-growth decomposition for belinostat at 10 *µ*M and 24 h. Columns show matched control cells, observed perturbed cells, velocity-only predictions, growth-only predictions and full velocity-plus-growth predictions. Grayscale indicates predicted growth weight. In this response, MCF7 was relatively less sensitive and was dominated by velocity-associated state change. K562 was dominated by growth-associated cell loss, whereas A549 required both components.

To assess U-Pert’s ability to recover drug-induced abundance changes, we compared the observed cell-abundance fold change with the mean predicted mass for each held-out condition. We evaluated gain over a no-change baseline in log2 fold-change space (Fig. 4b). Conditions were stratified by whether the observed fold change fell within the no-change band from 0.8 to 1.25. In conditions with measurable abundance shifts, U-Pert improved 79 of 81 held-out cases. The median error reduction was 0.466 log2 units (Wilcoxon *P* = 6.46 × 10^−15^). This result was not explained by a generic tendency to move predictions away from the no-change value of one. In conditions within the no-change band, only 14 of 43 cases improved over the no-change baseline. The median delta error was −0.069 (Wilcoxon *P* = 0.0074). Thus, U-Pert added value specifically when the perturbation required a change in population mass.

To assess how learning mass affected transcriptional-state prediction, we evaluated U-Pert across the same 124 held-out perturbation tuples. U-Pert retained strong state accuracy while adding an abundance readout (Fig. 4c,e). Relative to CellFlow, mean delta-expression correlation increased from 0.152 to 0.380, and mean gene-level R2 increased from 0.644 to 0.736. Latent energy distance decreased from 52.0 to 33.5, and latent Wasserstein distance decreased from 169.7 to 103.6. On these four displayed metrics, U-Pert also outperformed identity, chemCPA, Con-dOT and BioLord. Because supplementary state metrics showed less uniform separation among the strongest baselines, we do not interpret these results as uniform dominance across every transcriptional metric. Rather, the sciPlex3 benchmark supports a more specific conclusion: U-Pert recovered transcriptional responses while adding condition-specific abundance prediction.

To evaluate U-Pert’s generalization to unmeasured drug doses, we then applied the learned model to belinostat (PXD101) on a log-spaced dose grid across A549, K562 and MCF7 cells at 24 h (Fig. 4d). The predicted particle-level mass distributions followed the observed abundance trajectories across the measured dose points. This zero-shot interpolation converted dose into an explicit survival-like response. This expanded perturbation prediction beyond comparing expression states across doses.

Finally, the velocity-growth decomposition further separated distinct components of the drug response. We separated these components for belinostat at 10 *µ*M and 24 h (Fig. 4f), an HDAC inhibitor setting with cell-line-specific sensitivity. In this response, MCF7 was relatively less sensitive, and the observed effect was dominated by state displacement. Consistent with this biology, the velocity-only trajectory captured much of the MCF7 response. K562 showed the opposite pattern. The dominant effect was loss of cellular mass, and the growth-only trajectory moved closest to the observed treated population. A549 showed both effects, requiring both velocity and growth to match the observed response.

Together, these results show that in drug perturbation task, explicit mass modeling helps describe cellular responses that cannot be fully represented by expression-state shifts alone.

### U-Pert enables inverse drug screening for transcriptional and abundance phenotypes

We next asked whether the sciPlex3-trained U-Pert model could identify perturbations matching specified transcriptional or abundance phenotypes. For each design task, we kept the trained model fixed and optimized a continuous drug embedding to minimize the U-Pert objective for the desired response (Fig. 5a; Methods). The optimized embedding was then mapped to real compounds in an approximately 2,000-drug ChEMBL-derived library [43], and candidate compounds were re-evaluated with the same objective. Known pharmacology was used only after this step to assess the biological plausibility of the selected candidates.

**Fig. 5.**
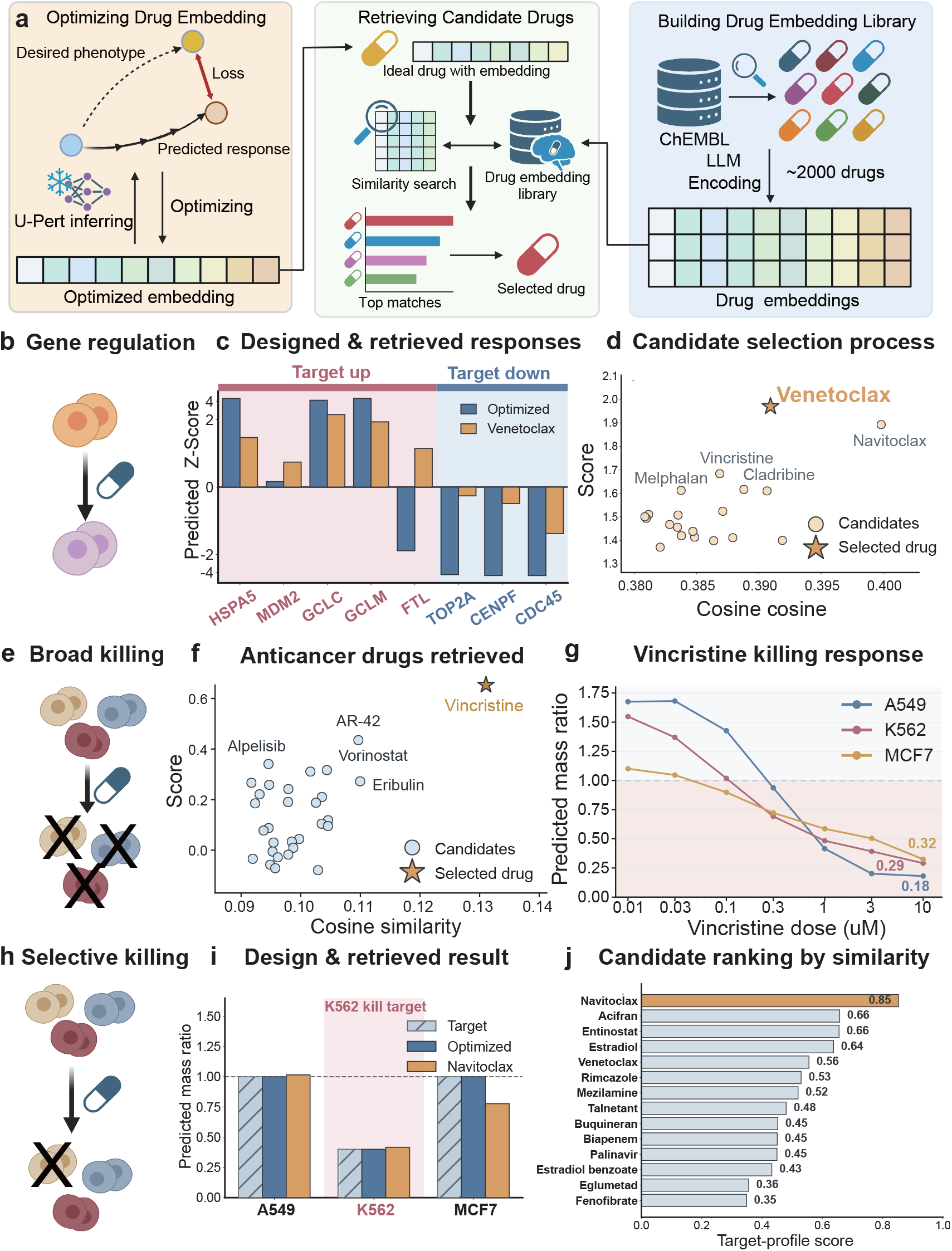
U-Pert enables inverse drug screening in sciPlex. **a**. Overview of the inverse cell-fate design and drug screening workflow. A continuous drug embedding is optimized through the trained U-Pert to match a desired perturbation phenotype, and the resulting optimized embedding is mapped to real compounds by similarity search in an LLM-derived drug-embedding library. The library was constructed by encoding approximately 2,000 ChEMBL-derived drug profiles with an LLM embedding model. **b**. Module-level design task in K562. **c**. Predicted module-gene responses for the optimized embedding and the selected real-drug candidate Venetoclax. **d**. Candidate mapping for the transcriptional module task. Each point represents a real drug evaluated by U-Pert after nearest-neighbor search in embedding space, and candidates are selected by task-specific validation score rather than cosine similarity alone. **e**. Broad-killing task, which aims to induce cell death across A549, K562 and MCF7. **f**. Candidate mapping for broad killing. Vincristine is selected as the top evaluated candidate. **g**. Dose-response validation for Vincristine across the three cell lines. **h**. K562-selective killing task, which targets reduced K562 survival while preserving A549 and MCF7 near baseline. **i**. Result of the K562-selective design. **j**. Candidate ranking for the selective-killing task by U-Pert target-profile score. Navitoclax ranks highest among evaluated candidates.

To design a transcriptional response, we targeted a K562 module-level response centered on cell-cycle regulation rather than isolated marker genes. The desired program increased a cellular-stress gene group, including HSPA5, MDM2, GCLC, GCLM and FTL, while suppressing a proliferative cell-cycle gene group marked by TOP2A, CENPF and CDC45 (Fig. 5b). These target modules reflect biological processes recurrently connected to unfolded-protein stress responses and cancer cell-cycle control [8, 39]. The optimized embedding strongly activated the target module, with a module score of 10.71. Mapping this designed embedding to the external compound library selected Venetoclax, whose predicted response preserved the intended module direction, increasing the cellular-stress gene group and suppressing the cell-cycle gene group while satisfying all specified up- and down-regulation directions (Fig. 5c,d). This candidate is biologically plausible: Venetoclax is a selective BCL-2 inhibitor with documented activity in BCL-2-dependent hematologic malignancy models, including chronic myeloid leukemia stem and progenitor contexts [44, 45].

To design an abundance phenotype, we then optimized a broad-killing objective intended to induce cell death across A549, K562 and MCF7 (Fig. 5e). The designed embedding reduced predicted survival across all three cell lines, and candidate mapping selected Vincristine as the top real compound, with predicted survival of 18%, 29% and 32% in A549, K562 and MCF7 at 10 *µ*M and a broad-killing score of 0.65 (Fig. 5f,g), which also shows a strong dosage dependence. Vincristine is a Vinca alkaloid microtubule-targeting chemotherapy, and microtubule disruption is a well-established anticancer mechanism that blocks mitotic progression and induces cell death [46]. Other high-scoring candidates in this screen (Alpelisib, Vorinostat, Eribulin, Arsenic Trioxide and Paclitaxel) also fell into recognizable oncology-related mechanisms, including PI3K inhibition, HDAC inhibition, microtubule targeting and pro-apoptotic leukemia therapy [47, 48]. The selected compound is therefore consistent with the broad-killing phenotype produced by the inverse design objective.

Finally, to evaluate U-Pert’s inverse-design capability for selective killing, we specified an abundance objective that reduced K562 survival while preserving the other cell lines. The target predicted survival values were 100%, 40% and 100% for A549, K562 and MCF7, respectively (Fig. 5h). The optimized embedding matched this target closely. After mapping the design to real compounds, Navitoclax ranked highest by target-profile score rather than by embedding similarity, reducing predicted K562 survival to 42% while keeping A549 near baseline at 102% and partially sparing MCF7 at 78% (Fig. 5i,j). This candidate is also mechanistically coherent: Navitoclax inhibits BCL-2, BCL-*x*_*L*_ and BCL-w, and BCL-family blockade engages mitochondrial apoptosis and can accelerate apoptosis during mitotic stress [49, 50]. Because K562 is a BCR-ABL-positive chronic myeloid leukemia-derived line, whereas A549 and MCF7 are non-leukemic solid-tumor cell lines, this leukemia-associated BCL-family vulnerability provides a plausible mechanism for the predicted K562-selective response.

Interestingly, the three tasks recovered pharmacologically coherent compounds: Venetoclax for the K562 transcriptional module, Vincristine for broad killing and Navitoclax for K562-selective killing. Notably, two K562-centered designs converged on BCL-family biology despite being driven by distinct objectives, consistent with studies showing that prosurvival BCL2-family programs support chronic myeloid leukemia stem/progenitor cells and that BCL-2/BCR-ABL co-targeting can eliminate these cells [45, 51]. This convergence suggests that U-Pert can prioritize biologically plausible perturbations for both transcriptional programs and cell abundance phenotypes.

### U-Pert preserves cytokine response geometry by reducing forced matching in PBMCs

We next considered a complementary regime in which large population-size changes are not the dominant signal. PBMC cytokine perturbations [52] provide such a setting, where responses are expected to be largely cell-type-specific, and the coupling of balanced transport may introduce implausible cross-cell-type matching. The Parse 10M PBMC dataset spans 12 donors, 90 cytokines and 16 immune cell types, providing a test of U-Pert’s ability to improve donor-held-out prediction by reducing forced matching and correcting response direction (Fig. 6a).

**Fig. 6.**
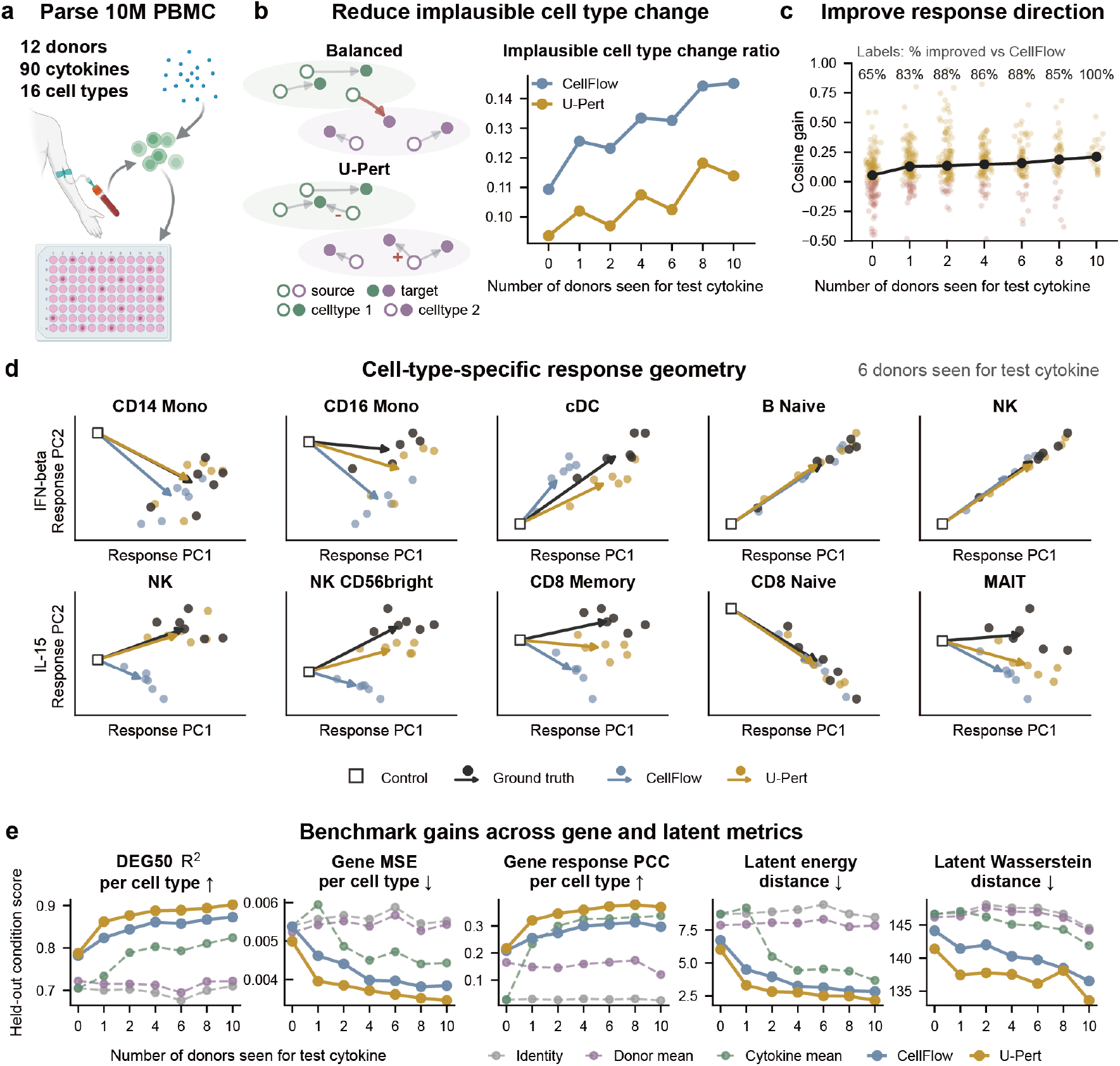
U-Pert preserves cytokine response geometry in PBMCs. **a**. The Parse 10M PBMC dataset contains short-term cytokine responses across 12 donors, 90 cytokines and 16 annotated immune cell types. **b**. Diagnostic for implausible cell-type reassignment under forced transport. Balanced transport-based prediction can encourage cross-cell-type matches, whereas U-Pert can modulate mass while retaining local response structure. Lower values indicate fewer assignments inconsistent with the annotated source and target cell types. **c**. Recovery of cytokine response direction across held-out donor settings. Response direction is quantified by cosine similarity between predicted and observed perturbation response vectors and aggregated at the condition level. Points show the median gain of U-Pert relative to CellFlow, and labels report the fraction of held-out cytokine-donor conditions improved by U-Pert. For each setting except 0 seen donors, the donor-generalization analysis was repeated three times. **d**. Cell-type-specific response geometry for IFN-*β* and IL-15. Each subpanel shows observed and predicted response vectors for biology-selected responder contexts at the 6-donor setting. IFN-*β* examples include CD14 monocytes, CD16 monocytes, conventional dendritic cells, naive B cells and NK cells; IL-15 examples include NK cells, CD56bright NK cells, memory CD8 T cells, naive CD8 T cells and MAIT cells. **e**. Summary benchmark across held-out perturbation metrics. U-Pert is compared with Identity, donor mean, cytokine mean and CellFlow.

To compare forced cross-cell-type matching between balanced and unbalanced formulations, we quantified the fraction of predicted mass assigned to cell-type changes inconsistent with the annotated source and target cell types. Balanced transport-based prediction enforces mass-preserving matching between control and treated distributions, which can encourage cross-cell-type assignments when the cytokine response is mainly a local transcriptional shift. By relaxing this constraint, U-Pert can avoid explaining every distributional difference through state transport across cell types. U-Pert reduced this wrong-change mass ratio relative to CellFlow across donor-generalization settings (Fig. 6b), from 0.109 to 0.094 when no donors were seen for the test cytokine, and from 0.131 to 0.118 at the 8-donor setting.

To link this reduction in forced matching to response geometry, we compared predicted and observed perturbation directions across held-out cytokine-donor conditions. The median cosine gain over CellFlow increased from 0.055 with no seen donors to 0.186 with 8 seen donors and 0.211 with 10 seen donors (Fig. 6c). At the 8-donor setting, U-Pert improved 85.4% of held-out conditions, with 48 condition-level summaries and a paired Wilcoxon test of *P* = 6.7 × 10^−7^. All non-zero seen-donor settings were repeated across three donor subsets.

To examine this directional correction in biologically relevant contexts, we analyzed known cytokine-responsive cell types treated with IFN-*β* and IL-15. For IFN-*β*, U-Pert better aligned predicted response vectors with observed responses in monocyte, dendritic, B cell and NK cell contexts. For IL-15, the same pattern appeared in NK, CD8 T cell and MAIT cell contexts (Fig. 6d). These cases show what the cosine metric captures: U-Pert keeps the response vector closer to the observed cell-type-specific cytokine direction, whereas balanced transport can distort the prediction direction through implausible assignments.

Notably, the gain in response geometry did not come at the expense of standard perturbation-prediction accuracy. At the 8-donor setting, U-Pert improved DEG50 R2 per cell type from 0.854 to 0.894, increased gene response Pearson correlation per cell type from 0.319 to 0.379, and reduced latent energy distance from 2.91 to 2.48 relative to CellFlow (Fig. 6e). Thus, U-Pert improved response direction while also achieving stronger gene-level and latent-distribution accuracy.

In summary, the PBMC analysis shows that unbalanced modeling remains useful even when large abundance shifts are not the main response. By relaxing mass-preserving matching, U-Pert reduced forced cross-cell-type assignments, improved response directions and retained strong gene-level and latent-distribution accuracy. In this setting, the unbalanced formulation of U-Pert achieved prediction robustness by preventing distributional mismatch from being converted into biologically implausible state transitions.

## Discussion

To capture the coupled effects of gene-expression change and cell-abundance variation from destructive and unpaired single-cell snapshots, we developed U-Pert, a mass-aware framework for single-cell perturbation modeling. By learning condition- and context-aware velocity and growth fields, U-Pert predicts responses to unseen perturbations and supports in silico prioritization of interventions for user-defined transcriptomic or population-level outcomes. The velocity field describes displacement in transcriptomic state space, whereas the growth field provides the abundance-associated information of expansion, depletion or selective recovery among profiled cells. Across genetic, chemical and cytokine perturbation settings, the endpoints measured by single-cell profiling were better represented as coupled molecular and population-level phenotypes than as purely transcriptomic states. These findings support treating recovered cell abundance as an explicit component of the perturbation response, while keeping separate the biological and technical factors that may contribute to this signal.

Existing single-cell perturbation models have achieved substantial progress in counterfactual expression prediction, response mapping and generative modeling [13–15, 21, 23, 24, 29]. Their main object, however, is usually the perturbed expression distribution. When the recovered endpoint population differs in size or composition, balanced formulations may absorb depletion, expansion or selective survival into apparent transcriptomic movement. Related unbalanced and growth-aware approaches in trajectory inference have shown that proliferation, apoptosis and changes in population size can be incorporated into dynamical models of single-cell data [41, 42, 53–56], where unbalanced transport and growth-aware models reconstruct cellular dynamics while accounting for unbalance sampling, proliferation, apoptosis and shifts in population size. These methods show that population-size changes can be treated as part of the dynamical process rather than as a nuisance variable. However, they primarily address autonomous temporal progression, whereas perturbation biology requires models that are conditioned on external genetic or pharmacological interventions, biological context and design objectives. U-Pert brings this unbalanced perspective into single-cell perturbation modeling by coupling transcriptomic state transitions with perturbation-induced proliferation and apoptosis. This separation provides a more interpretable description of perturbation responses, allowing drug effects to be decomposed into state-transition and survival components, and cytokine responses to be modeled without forcing implausible cross-cell-type matching. It also creates a route to inverse design, because desired outcomes can be specified at the level of gene-expression programs, cell-type compositions or abundance phenotypes.

While this study demonstrates the potential of unbalanced perturbation modeling, several limitations remain. First, observed cell-count changes can reflect both biological effects and technical factors, including sampling depth, capture efficiency and batch variation. More carefully controlled experimental designs will be important for separating true proliferation or apoptosis from technical sampling noise. Second, because U-Pert learns dynamics from unpaired snapshots, the inferred trajectories should be interpreted as plausible transport paths rather than directly observed single-cell lineages. Future integration with lineage tracing [57], live-cell imaging [58] or densely sampled time-resolved perturbation measurements [24] could further constrain the learned dynamics. Third, the current inverse-design workflow relies on learned perturbation or drug embeddings and candidate retrieval in an embedding library; therefore, experimental validation is required before designed perturbations can be interpreted as causal or therapeutically actionable interventions.

Future extensions could broaden the biological scope of unbalanced perturbation modeling. Spatial perturbation technologies now make it possible to measure genetic perturbations together with spatial transcriptomic or imaging measurements in native tissue contexts [59–63]. Extending U-Pert to these data would allow perturbation responses to be modeled across transcriptomic state, population mass, physical location and local cell–cell communication. Such models could distinguish how perturbations alter the targeted cells themselves from how they reshape neighboring cells through ligand–receptor signaling, niche remodeling and collective tissue-level responses [64]. Multimodal measurements, perturbation time courses and prospective validation of designed interventions would also further clarify whether the mass-aware virtual-cell models can support experimental decision-making.

Overall, U-Pert provides a unified unbalanced generative framework for learning perturbation-induced cell-state and cell-abundance dynamics from unpaired unbalanced single-cell snapshots. By combining forward simulation, mechanistic interpretation and inverse design, U-Pert extends virtual-cell modeling from predicting transcriptional responses to designing perturbations that control both molecular programs and population outcomes. More broadly, our results support an unbalanced view of perturbation biology, in which virtual-cell models incorporate population-level changes alongside transcriptional state to predict and design cell fate.

## Methods

### Learning cell-state and abundance dynamics from unpaired snapshots

The dynamic unbalanced optimal transport described by the Wasserstein–Fisher–Rao (WFR) metric extends classical OT by coupling displacement with changes in mass, providing a geometry for modeling cell-state transitions together with proliferation, apoptosis or selection[65–67]. In our setting, *µ*_0_ and *µ*_1_ denote finite measures over the single-cell state space *X*, representing the control and perturbed cell populations. Unlike balanced OT, which preserves total mass, WFR allows the total mass of the population to increase or decrease along the path. The dynamic WFR distance is defined as[65, 66]:

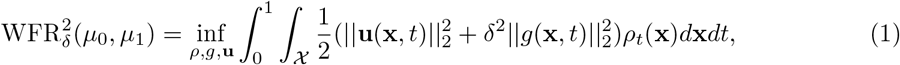

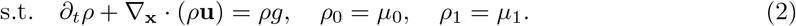

Here, **u**(**x**, *t*) is the velocity field that transports transcriptomic states, *g*(**x**, *t*) is the growth rate that changes cell mass, *ρ*_*t*_(**x**) is the time-dependent cell distribution, and *δ* balances transport and birth-death penalties. Positive values of *g* locally increase mass, whereas negative values decrease mass.

To connect this dynamic formulation to flow matching, we use the equivalent static WFR semi-coupling view[65, 66]. A semi-coupling (*γ*_0_, *γ*_1_) consists of two non-negative measures on *X* ^2^. The first measure *γ*_0_ specifies how much source mass at **x** is assigned to a source–target pair (**x, y**), whereas *γ*_1_ specifies the corresponding target mass at **y**. Thus, each pair can carry different source and target masses, recording both movement and birth-death effects. The static WFR formulation is:

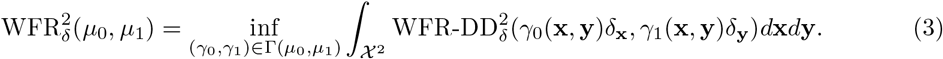

Here, WFR-DD_*δ*_ denotes the WFR cost between two weighted Dirac measures, and Γ(*µ*_0_, *µ*_1_) is the admissible set of semi-couplings,

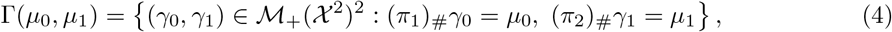

where *π*_1_(**x, y**) = **x** and *π*_2_(**x, y**) = **y** are the coordinate projections. In other words, the source marginal of *γ*_0_ recovers the control population, and the target marginal of *γ*_1_ recovers the perturbed population. In practice, the semi-coupling is computed through an equivalent optimal entropy-transport formulation [42, 65, 66].

WFR Flow Matching (WFR-FM)[42] learns a time-dependent vector field **v**_***θ***_(**x**, *t*) and growth rate *g*_***ϕ***_(**x**, *t*) from the semi-coupled source–target pairs. For a sampled boundary pair **z** = (**x**_0_, **x**_1_) from *γ*_0_, WFR-FM defines a conditional path 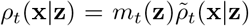. Here, 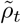 is the normalized conditional distribution of cell states along the path, and *m*_*t*_(**z**) is the WFR mass associated with that path at time *t*. The WFR geodesic provides target fields **u**_*t*_(**x**|**z**) for transcriptomic movement and *g*_*t*_(**x** | **z**) for mass change. The neural networks are then optimized with the conditional unbalanced flow matching objective:

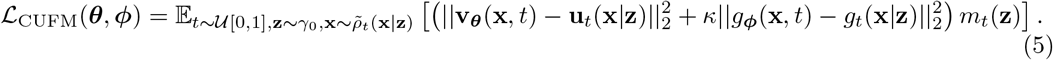

The mass weight *m*_*t*_(**z**) makes the regression objective sensitive to birth-death dynamics rather than only to transcriptomic displacement. The closed-form construction of the conditional path and target fields follows from the WFR geodesic [42, 65, 66].

### Learning condition- and context-aware perturbation responses

To apply WFR-FM to single-cell perturbation modeling, U-Pert distinguishes between perturbation conditions, biological contexts and the matching strata used to construct WFR couplings. The perturbation condition, denoted by cond, specifies the intervention and intervention-related variables, such as a genetic knockout, drug identity, dose or treatment duration. The biological context, denoted by cont, specifies the basal or experimental background in which the perturbation acts, such as donor identity, cell line or cell type.

In practice, WFR couplings are constructed within dataset-specific matching strata. Each observed stratum *s* is associated with a perturbation condition cond_*s*_ and, when available, the context covariates used to match control and perturbed cells. For example, when control and perturbed cells from the same donor or cell line are available, the stratum can include that donor or cell-line label; otherwise, the coupling may be constructed at a coarser condition level while the context is provided as an additional neural-network input. This design allows U-Pert to use matched controls whenever the experimental design supports them, while still sharing information across contexts through the learned condition- and context-aware fields.

For each observed stratum *s*, let 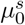 and 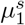 denote the corresponding matched control and perturbed distributions. U-Pert solves the static WFR problem between 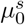 and 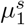 to obtain stratum-specific semi-couplings 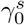 and 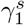. The trainable velocity **v**_***θ***_(**x**, *t*, cond, cont) and growth rate *g*_***ϕ***_(**x**, *t*, cond, cont) are conditioned on both the perturbation condition and the biological context. The training objective is

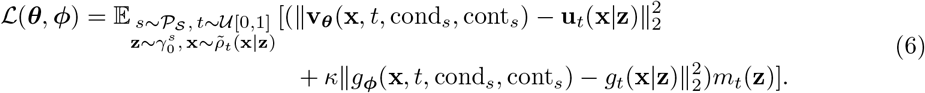

Here, **z** denotes a WFR source-target pair sampled from the semi-coupling, *m*_*t*_(**z**) is the corresponding WFR path mass, and cond_*s*_ and cont_*s*_ are the condition and context labels provided to the neural networks for stratum *s*. This objective allows U-Pert to learn how the same control population can undergo different transcriptomic transitions and abundance changes under different interventions and biological backgrounds.

For prediction, U-Pert uses multi-step integration to simulate the coupled state and growth dynamics under a specified perturbation condition and biological context:

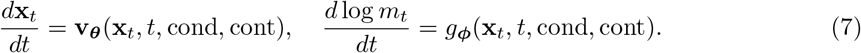

The same condition- and context-aware framework can also be combined with a mean-flow parameterization for one-step terminal prediction [56, 68]. In this variant, the network directly predicts terminal displacement and terminal log-mass change, avoiding backpropagation through the ODE solver. We therefore treat it as an optional acceleration strategy, particularly useful for large-scale gradient-based inverse design.

### Preprocessing and implementation of U-Pert

#### Cell embedding

Single-cell transcriptional states were preprocessed before U-Pert training. For all real biological datasets, count matrices were normalized, log-transformed, filtered to retain highly variable genes (HVGs), and projected to a 100-dimensional PCA representation. The PCA model was fitted using control cells and training perturbation conditions. U-Pert learned transport and growth in this PCA space. Predicted PCA states were subsequently reconstructed in the HVG expression space for gene-level analyses.

#### Condition embedding

Each perturbation was represented by a fixed external embedding. The embedding source depended on the perturbation modality and could include textual descriptions, molecular features or protein annotations. In the real-data applications, compound and cytokine descriptions were embedded before model training. U-Pert mapped each raw perturbation embedding through a multilayer condition encoder consisting of layer normalization, two feed-forward hidden layers and a final linear projection. The hidden dimension was 512, and the output condition representation was 256-dimensional in both applications.

#### Context embedding

Experimental and biological contexts were encoded separately from perturbation identity according to a dataset-specific rulebook. Each prediction target was represented as a condition tuple, and the rulebook specified which covariates were used for control matching and model conditioning. Categorical covariates were encoded by trainable embedding layers, whereas continuous covariates were transformed as specified by the rulebook and encoded by two-layer multilayer perceptrons. During training and inference, control cells were selected with the same matching covariates used to define the biological context.

#### Model architecture

U-Pert comprised two conditional output networks: a velocity network for transcriptomic transport and a growth network for particle mass dynamics. The networks had separate parameters but shared the same conditioning inputs. For each sampled particle, the model encoded the perturbation, WFR interpolation time and context covariates, and then concatenated these representations to condition the output networks. WFR time was represented by sinusoidal features followed by a feed-forward time encoder. The processed time representation was projected to 128 dimensions.

The velocity and growth networks used FiLM-conditioned residual multilayer perceptron blocks [69, 70]. Each block first mapped the particle state into a hidden representation, normalized the hidden state and generated FiLM scale, shift and gate parameters from the concatenated conditioning vector. The block then applied a two-layer feed-forward transformation and added the gated output to the residual state. The final FiLM projection was initialized to zero, making each residual block close to an identity map at initialization. The velocity network output a 100-dimensional vector field in PCA space, whereas the growth network output one scalar growth rate per particle.

#### Implementation

U-Pert was implemented in Python with PyTorch. Unbalanced transport plans were computed with POT [71]. Data preprocessing, AnnData storage and gene-expression analyses used the scverse ecosystem [72], including anndata [73] and scanpy [74].

### Real-data perturbation benchmarks

For real biological data, each prediction target was defined by a perturbation identity together with a biological or experimental context. U-Pert used matched control cells as the starting population for each target context and predicted both terminal cell states and particle masses. State-level predictions were evaluated in the shared PCA and HVG expression spaces, whereas mass predictions were used to assess abundance changes when cell-count information was available.

In the genetic perturbation benchmarks, we evaluated unseen-condition prediction on Norman, Adamson and Replogle K562 Perturb-seq datasets [8, 9, 11]. Prediction targets were genetic perturbation conditions, and matched control cells were used as source populations. We compared U-Pert with CellFlow, scGPT, GEARS, perturbation-mean and zero-change baselines for expression and distributional metrics, and with a supervised MLP baseline for condition-level abundance-ratio prediction.

In sciPlex3 datasets [7], we evaluated chemical perturbation responses across 188 compounds, three cancer cell lines, four dose levels and two time points. A prediction target was defined by a compound-cell line-dose-time tuple. We used a held-out-drug split to test generalization to unseen compounds and evaluated both transcriptional-state recovery and drug-induced cell-abundance change. Using the trained response field, we further performed inverse drug screening for user-defined gene-expression and abundance objectives.

In the PBMC cytokine-perturbation datasets [52], we evaluated responses across 12 donors and 90 cytokines profiled after 24 h. A prediction target was defined by a cytokine-donor pair. We used held-out-cytokine and seen-donor generalization settings with donor-matched control cells, and assessed whether U-Pert preserved cytokine response direction while reducing forced cross-cell-type transport.

### Evaluation Metrics

To evaluate predicted perturbation responses, we adopted six population-average and population-distribution metrics from recent single-cell perturbation benchmarks [38]: population-average MSE, population-average E-distance, population-average PCC-delta, population-distribution Wasserstein distance, population-distribution KL-divergence and population-distribution Common-DEGs. To explicitly evaluate cellular birth-death dynamics and cell-fate shifts, we additionally defined the following U-Pert-specific abundance and composition metrics.

#### Cell Type Proportion Metrics

Traditional metrics evaluate transcriptomic states but ignore population expansions or contractions. To evaluate how well the model captures unbalanced cell-fate dynamics, we evaluate the predicted post-perturbation cell type proportions against the ground truth. To assign cell types to the predicted perturbed cells in a non-parametric manner, we determine the identity of each predicted cell via a majority vote from its *k*-nearest neighbors (kNN) in the true perturbed cell pool.

Let **p** ∈ Δ^*K*−1^ and 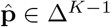 denote the true and predicted cell type proportion distributions across *K* defined cell types. We quantify the compositional accuracy using the Mean Squared Error (MSE_prop_) and the Kullback-Leibler (KL) divergence (KL_prop_):

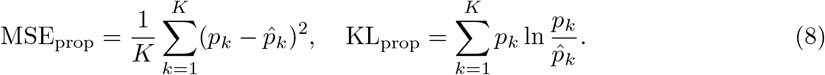

These custom metrics explicitly reflect the fidelity of the modeled birth-death dynamics under perturbation conditions.

#### Mass fold-change and abundance gain

For models that predict particle masses, condition-level abundance change was evaluated from the total predicted mass. If *M*_0_ and *M*_1_ denote the observed control and perturbed population masses, and 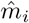 denotes the predicted terminal mass of control particle *i*, the observed and predicted fold changes are:

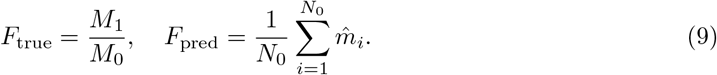

Mass error was measured as (*F*_true_ − *F*_pred_)^2^. In the sciPlex3 abundance analysis, gain over a no-change prediction was measured in log2 fold-change space:

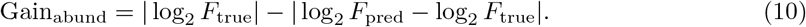

Positive values indicate lower abundance error than a no-change baseline.

#### Cell-type transition error

For the PBMC analysis, we quantified implausible source-to-predicted cell-type transitions. Let *s*_*i*_ and *ĉ*_*i*_ denote the source and predicted cell-type labels for particle *i*, and let *A*(*s*_*i*_) denote the allowed target cell types for source type *s*_*i*_. The mass fraction assigned to forbidden transitions was:

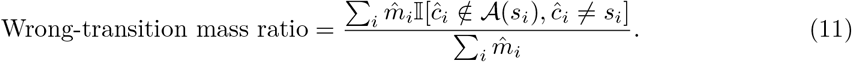

We also reported the same numerator divided by the total mass assigned to changed cell-type labels.

#### PBMC response-direction cosine gain

To evaluate donor- and cell-type-specific cytokine response direction, we computed cosine similarity between observed and predicted response vectors in PCA space. For a cytokine, donor and cell-type context, let:

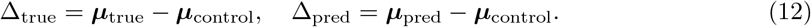

The response-direction score is:

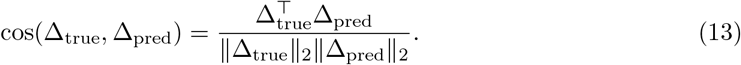

U-Pert gain was defined as the U-Pert cosine similarity minus the CellFlow cosine similarity.

### Discovery and Enrichment Analysis of Driver Genes

Beyond predicting the macroscopic terminal states of perturbed cell populations, understanding the underlying molecular mechanisms that drive these transitions is essential for biological discovery [6]. Our generative framework parameterizes the continuous dynamics of transcriptomic state and cellular mass, providing a differentiable foundation for extracting model-inferred gene regulatory relationships and growth-associated effects [53, 54, 75].

Given a specific perturbation condition cond and cellular context cont, the velocity field **v**_***θ***_(**x**, *t*, cond, cont) dictates the instantaneous transcriptomic rate of change in the learned expression representation. For real biological datasets, U-Pert is trained in a low-dimensional PCA space. Let **y** ∈ ℝ^*d*^ denote the highly variable gene expression vector and **x** ∈ ℝ^*r*^ denote its PCA representation, where *d* is the number of genes and *r* is the number of retained principal components.

We write the linear PCA encoder and decoder as

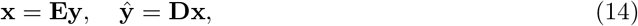

where centering, scaling and intercept terms are omitted for notation simplicity.

We first compute the Jacobian matrix of the velocity field in PCA space:

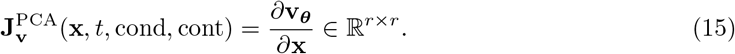

To obtain a gene-space regulatory sensitivity matrix, we project this PCA-space Jacobian back to gene space by the chain rule:

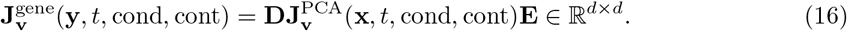

In this formulation, the element 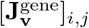 represents the model-inferred instantaneous regulatory effect of gene *j* on the expression velocity of gene *i*, after projection through the PCA encoder and decoder.

Simultaneously, the growth rate function *g*_***ϕ***_(**x**, *t*, cond, cont) models condition-dependent changes in cellular mass. We compute the gradient of the growth rate in PCA space,

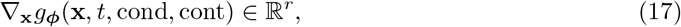

and project it back to gene space using the PCA encoder:

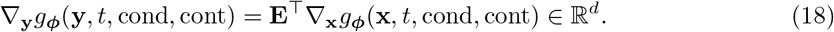

This gene-level growth sensitivity score quantifies how variation in each gene is associated with local increases or decreases in the predicted growth rate. Depending on the experimental setting, these growth-associated effects may reflect proliferation, apoptosis, selective recovery or other sources of abundance change.

Because the underlying neural networks are explicitly conditioned, the inferred gene regulatory networks 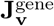 and growth sensitivity landscapes ∇_**y**_*g*_***ϕ***_ are condition- and context-dependent. They are not static global properties; rather, they can rewire in response to different perturbation interventions or cellular backgrounds. By contrasting the projected Jacobians and growth gradients across different cond and cont embeddings, we can systematically probe how specific drugs or genetic knockouts alter gene circuitry and abundance-associated pathways.

Genes exhibiting the highest magnitude of change in their regulatory outgoing weights, measured by large column norms in 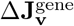, or growth influence, measured by large absolute values in Δ ∇_**y**_*g*_***ϕ***_, are identified as putative driver genes for that specific condition. Finally, to translate these mathematical sensitivities into actionable biological insights, the identified sets of critical driver genes are subjected to downstream enrichment analyses, mapping them to known biological pathways, gene ontologies and disease-associated mechanisms [76, 77].

### Inverse Design of Differential Expression Genes Task

The ultimate goal of modeling single-cell perturbation dynamics is to enable *in silico* inverse design: computationally identifying a perturbation condition cond that drives a cell population toward a desired biological state. In many biological applications, this desired state is characterized by the differential expression of specific marker genes. Formally, we define a set of target genes to be up-regulated, denoted as *S*_up_, and a set of target genes to be down-regulated, denoted as *S*_down_. The inverse design task is to optimize the condition embedding cond to induce the desired expression changes in these target gene sets.

Let Φ_cond,cont_ denote the differentiable U-Pert forward operator under a given perturbation condition and biological context. This operator maps an initial control cell, together with its initial particle mass, to a predicted perturbed state and predicted terminal mass:

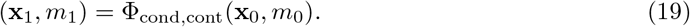

The operator Φ_cond,cont_ can be implemented either by multi-step integration of the learned WFR-FM dynamics or by the optional one-step mean-flow variant [56]. In the inverse design objective below, only the differentiable input-output mapping of Φ_cond,cont_ is required.

Given a set of *N* control cells 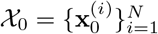, each initialized with mass 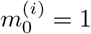, we simulate their response to a proposed condition cond:

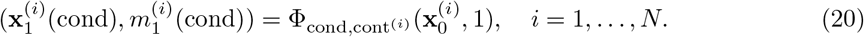

For DEG-based design, the predicted masses are normalized and used as sample weights for the predicted perturbed population:

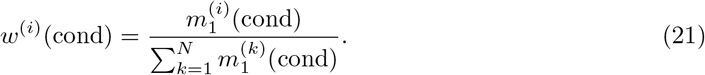

This normalization makes the DEG objective focus on the mass-weighted expression profile of the predicted recovered population. Total abundance or survival objectives are handled separately in the abundance-design tasks.

Let 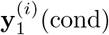 denote the predicted gene-expression vector used for evaluating target genes, obtained from 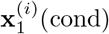 (cond) by PCA reconstruction when the model is trained in PCA space. Let 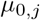 and 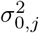 denote the empirical mean and variance of gene *j* in the control population. For the predicted perturbed population, we compute the mass-weighted mean and variance:

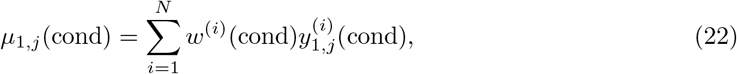

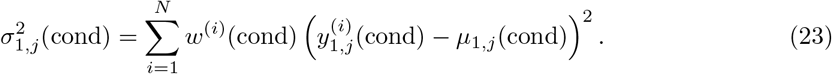

We use a t-statistic-inspired standardized expression contrast to quantify the direction and magnitude of the predicted expression change for each target gene *j*:

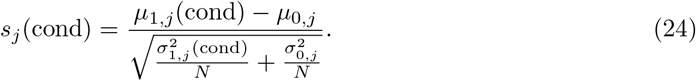

The design objective maximizes *s*_*j*_(cond) for genes that should be up-regulated and minimizes *s*_*j*_(cond) for genes that should be down-regulated:

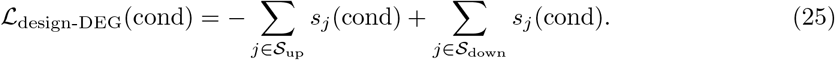

The optimized perturbation condition is then obtained by

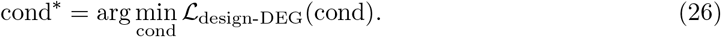

Because the forward operator Φ_cond,cont_ is differentiable, this optimization can be performed by backpropagating the loss through the trained U-Pert model while keeping model parameters fixed and updating only the continuous condition embedding. For discrete perturbations, such as drug compounds or target genes, the optimized embedding is subsequently mapped back to real candidates in a predefined perturbation-embedding library and the retrieved candidates are re-evaluated by forward U-Pert inference.

### Inverse Design of Cell Type Proportion/Quantity Task

Beyond targeting specific gene expression profiles, a major goal in regenerative medicine, cell therapy, and precision oncology is to discover perturbation interventions that can control cell population dynamics. Practical applications include directing stem cells to differentiate into a desired lineage, selectively expanding a specific immune subpopulation, or designing targeted drugs that eradicate cancer cells while preserving healthy tissues. To achieve these macroscopic objectives, modeling only transcriptomic transitions of individual cells is insufficient; one must also capture the expansion and depletion of the populations themselves. Therefore, explicitly modeling growth dynamics is important for this task. We formalize this objective as the inverse design of cell type proportions, cell type quantities, or abundance ratios of fixed biological groups.

As in the differential expression task, we use the differentiable forward operator Φ_cond,cont_ to simulate the perturbation response of *N* control cells. Given control cells 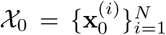, each initialized with mass 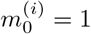, the predicted perturbed states and terminal masses are

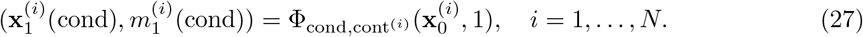

We consider two related settings. In the first setting, the target population label is state-dependent and may change after perturbation, as in cell fate conversion or differentiation. To enable gradient-based optimization of the condition embedding cond, the assignment of predicted perturbed cells to specific cell types must be differentiable. Therefore, non-differentiable assignment methods, such as *k*-nearest neighbors (kNN) or hard clustering, cannot be used. Instead, we pre-train a continuous neural network classifier *C*(**x**) ∈ [0, 1]^*K*^, where *K* is the total number of defined cell types. For any given cell state **x**, the output *C*_*k*_(**x**) represents the predicted probability that the cell belongs to cell type *k*, subject to 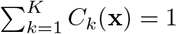.

The unbalanced formulation provides a natural way to optimize not only relative cell type proportions, but also the absolute quantity of cells assigned to each type. The differentiable predicted quantity *Q*_*k*_(cond) for cell type *k* is computed by summing the predicted masses weighted by the classifier assignment probabilities:

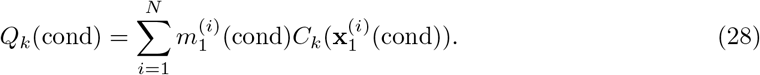

Similarly, the predicted relative proportion *P*_*k*_(cond) for cell type *k* is obtained by normalizing this cell type quantity by the total predicted mass:

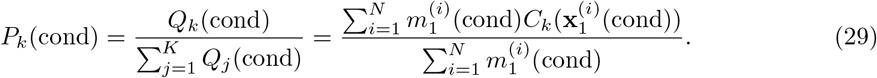

Given a user-defined target quantity profile 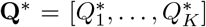 or a target proportion profile 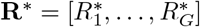, we formulate the state-dependent cell type design objective using mean squared error:

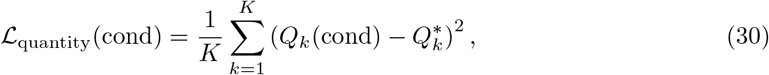

and

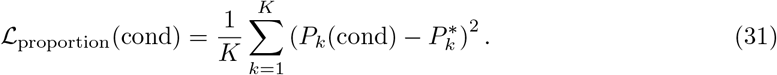

In the second setting, the target group label is fixed and does not change with the transcriptomic state. Examples include cell line, donor, sample group or other predefined biological contexts. This setting is used, for example, when designing a drug that selectively kills one cell line while preserving others. Let *a*^(*i*)^ ∈ {1, …, *G*} denote the fixed group label of control cell *i*, where *G* is the number of fixed groups. The initial mass of group *g* is

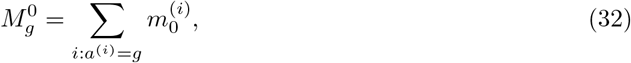

and the predicted terminal mass of group *g* under condition cond is

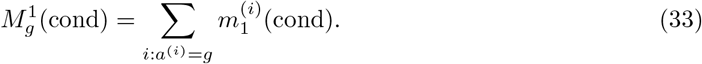

The predicted abundance ratio for group *g* is then defined as

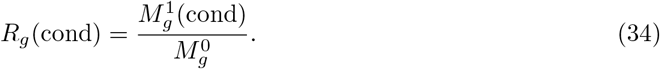

Given a target abundance-ratio profile 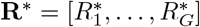, the fixed-group abundance design loss is

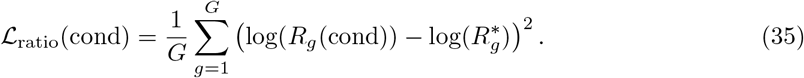

For example, a selective-killing objective can be specified by setting a low target ratio for the cell line to be depleted and target ratios near one for the cell lines to be preserved.

The optimized perturbation condition is identified by minimizing the corresponding target loss:

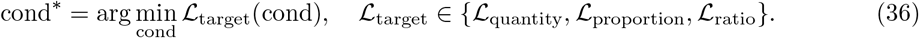

Because the classifier *C*(·), when used, and the forward operator Φ_cond,cont_ are differentiable, the condition embedding can be optimized by backpropagating the target loss through the frozen classifier and the trained U-Pert model. For fixed-group abundance-ratio objectives, no classifier is required because the group label is fixed and carried by each particle during forward simulation. For discrete perturbations, such as drug compounds or target genes, the optimized continuous embedding is subsequently mapped back to real candidates in a predefined perturbation-embedding library and the retrieved candidates are re-evaluated by forward U-Pert inference.

### Simulations of perturbed gene regulatory networks

To evaluate model performance under diverse perturbations, we simulated single-cell gene expression dynamics using a three-gene regulatory network. The dynamics of the system are governed by the following set of stochastic differential equations (SDEs):

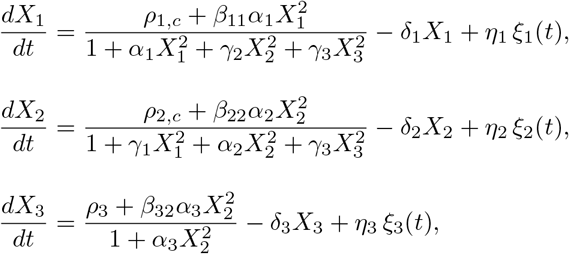

where *X*_*i*_(*t*) denotes the mRNA expression level of gene *i*. The model implements a toggle switch between *X*_1_ and *X*_2_ (mutual inhibition and self-activation), with additional regulation by *X*_3_ and an external signal *β*. The equations are parameterized by basal transcription rates (*ρ*_*i,c*_), self-activation strengths (*α*_*i*_), inhibitory interactions (*γ*_*i*_), degradation rates (*δ*_*i*_), and stochastic noise terms (*η*_*i*_*ξ*_*i*_(*t*)). Probabilistic cell death is incorporated by assigning each cell a death probability over a time step *dt*:

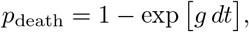

where the instantaneous growth rate *g* depends on gene expression:

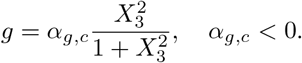

Here, *α*_*g,c*_ is a condition-specific parameter, with more negative values corresponding to stronger apoptosis.

For each perturbation condition, an initial population of up to 10,000 cells was sampled from the control distribution. Each condition *c* is defined by a triplet of parameters (*ρ*_1,*c*_, *ρ*_2,*c*_, *α*_*g,c*_), randomly drawn from predefined ranges. A total of 200 perturbation conditions were generated, with 100 assigned to the training set and 100 reserved for evaluation. Each condition was simulated independently in parallel, producing final single-cell populations that capture the combined effects of network regulation, stochastic fluctuations, and condition-specific survival.

To define cell-type composition in the simulated system, we assigned each cell to one of four expression-defined states based on the relative dominance of the three gene expression levels. For each cell, raw expression values (*X*_1_, *X*_2_, *X*_3_) were first transformed to positive values using a softplus function and normalized to obtain relative expression fractions:

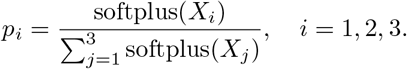

We then quantified whether a cell showed a dominant gene-expression program by computing the difference between the largest and second-largest relative expression fractions. Cells with a sufficiently dominant gene were assigned to Type 1, Type 2, or Type 3 according to the gene with the highest relative expression. Cells without a clearly dominant gene-expression program were assigned to Type 4, representing an intermediate or non-dominant state.

For differentiable optimization of cell-type composition during inverse design, we used a soft assignment scheme. Dominance was controlled by a sigmoid gate with margin 0.10 and temperature 0.03, and gene-specific class probabilities were computed using a softmax with temperature 0.03. The resulting four-dimensional soft cell-type vector represents probabilities for Type 1, Type 2, Type 3, and Type 4. Population-level cell-type proportions were then computed as the normalized weighted average of these soft assignments, using the predicted cell weights from U-Pert.

### sciPlex drug inverse design and candidate retrieval

For the sciPlex inverse-design analysis, we used the trained sciPlex U-Pert model as a fixed differentiable forward operator and optimized only the continuous drug embedding. Cell-line context, dose and time covariates were fixed according to the target task. We considered three objectives: a K562 transcriptional gene-regulation objective, a broad-killing objective that minimized predicted mass across A549, K562 and MCF7, and a selective-killing objective that matched target mass ratios across the three cell lines.

Candidate drugs were selected from an external merged library of 2,009 ChEMBL-derived and curated clinical candidate records [43]. For each drug, an LLM first generated a structured pharmacology description covering targets, mechanisms, pathways, expected transcriptional programs and cell-line-specific effects, and this text was embedded with a text-embedding model. In total, 1,985 candidates had valid embeddings. After optimizing a continuous embedding, we retrieved nearest real-drug candidates by cosine similarity and then re-ran U-Pert forward inference for each candidate under the same task settings. Final compounds were selected by task-specific validation scores from this re-inference step, not by cosine similarity alone.

## Data Availability

All real datasets used in this paper are publicly available. The Norman [11], Adamson [8] and Replogle K562 [9] Perturb-seq datasets used for the genetic perturbation benchmarks are publicly available and were accessed through the GEARS data utilities [18]. The sci-Plex3 chemical perturbation dataset was obtained from the public sci-Plex data release [7]. The PBMC cytokine-perturbation dataset [52] was obtained from the publicly available Parse Biosciences 10 Million Human PBMCs in a Single Experiment resource, available through the Parse Biosciences data portal at https://www.parsebiosciences.com/datasets/10-million-human-pbmcs-in-a-single-experiment/. The simulated datasets generated in this study will be released upon acceptance of the manuscript.

## Code Availability

The code of U-Pert will be made publicly available upon acceptance of the manuscript.

## Acknowledgments

P.Z. acknowledges support from the National Natural Science Foundation of China under grants 12288101, 8206100646, and T2321001, the Fundamental and Interdisciplinary Disciplines Break-through Plan of the Ministry of Education of China under grant JYB2025XDXM502, and the Fundamental Research Funds for the Central Universities. We also acknowledge the support from the High-performance Computing Platform of Peking University for computation.

## Contribution

Q.P. and P.Z. conceived the study. Q.P., Y.W., and J.L. developed and implemented the computational method. Q.P., Y.W., J.L., X.W., and Y.X. performed the data analysis and wrote the original draft. All authors reviewed and edited the manuscript. P.Z. supervised the project.

## References

[1] MacArthur, B. D., Ma’ayan, A. & Lemischka, I. R. Systems biology of stem cell fate and cellular reprogramming. Nature Reviews Molecular Cell Biology 10, 672–681 (2009).

[2] Moris, N., Pina, C. & Arias, A. M. Transition states and cell fate decisions in epigenetic landscapes. Nature Reviews Genetics 17, 693–703 (2016).

[3] Perrimon, N., Pitsouli, C. & Shilo, B.-Z. Signaling mechanisms controlling cell fate and embryonic patterning. Cold Spring Harbor Perspectives in Biology 4, a005975 (2012).

[4] Bunne, C. et al. How to build the virtual cell with artificial intelligence: Priorities and opportunities. Cell 187, 7045–7063 (2024).

[5] Viñas Torné, R. et al. Systema: a framework for evaluating genetic perturbation response prediction beyond systematic variation. Nature Biotechnology 1–10 (2025).

[6] Dixit, A. et al. Perturb-Seq: dissecting molecular circuits with scalable single-cell RNA profiling of pooled genetic screens. Cell 167, 1853–1866 (2016).

[7] Srivatsan, S. R. et al. Massively multiplex chemical transcriptomics at single-cell resolution.Science 367, 45–51 (2020).

[8] Adamson, B. et al. A multiplexed single-cell CRISPR screening platform enables systematic dissection of the unfolded protein response. Cell 167, 1867–1882 (2016).

[9] Replogle, J. M. et al. Mapping information-rich genotype-phenotype landscapes with genomescale perturb-seq. Cell 185, 2559–2575 (2022).

[10] Datlinger, P. et al. Pooled CRISPR screening with single-cell transcriptome readout. Nature Methods 14, 297–301 (2017).

[11] Norman, T. M. et al. Exploring genetic interaction manifolds constructed from rich single-cell phenotypes. Science 365, 786–793 (2019).

[12] Maan, H. et al. Characterizing the impacts of dataset imbalance on single-cell data integration.Nature biotechnology 42, 1899–1908 (2024).

[13] Lotfollahi, M., Wolf, F. A. & Theis, F. J. scGen predicts single-cell perturbation responses.Nature Methods 16, 715–721 (2019).

[14] Lotfollahi, M., Naghipourfar, M., Theis, F. J. & Wolf, F. A. Conditional out-of-distribution generation for unpaired data using transfer VAE. Bioinformatics 36, i610–i617 (2020).

[15] Lotfollahi, M. et al. Predicting cellular responses to complex perturbations in high-throughput screens. Molecular Systems Biology 19, MSB202211517 (2023).

[16] Hetzel, L. et al. Predicting cellular responses to novel drug perturbations at a single-cell resolution. Advances in Neural Information Processing Systems 35, 26711–26722 (2022).

[17] Aliee, H. et al. inVAE: Conditionally invariant representation learning for generating multivariate single-cell reference maps. bioRxiv 2024–12 (2024).

[18] Roohani, Y., Huang, K. & Leskovec, J. Predicting transcriptional outcomes of novel multigene perturbations with gears. Nature Biotechnology 42, 927–935 (2024).

[19] Kamimoto, K. et al. Dissecting cell identity via network inference and in silico gene perturbation. Nature 614, 742–751 (2023).

[20] Wang, T. et al. Cellnavi predicts genes directing cellular transitions by learning a gene graph-enhanced cell state manifold. Nature Cell Biology 27, 1863–1874 (2025).

[21] Bunne, C. et al. Learning single-cell perturbation responses using neural optimal transport.Nature Methods 20, 1759–1768 (2023).

[22] Dong, M. et al. Causal identification of single-cell experimental perturbation effects with CINEMA-OT. Nature Methods 20, 1769–1779 (2023).

[23] Klein, D. et al. CellFlow enables generative single-cell phenotype modeling with flow matching.bioRxiv 2025–04 (2025).

[24] Rohbeck, M. et al. Modeling complex system dynamics with flow matching across time and conditions. The Thirteenth International Conference on Learning Representations (2025).

[25] Hao, M. et al. Large-scale foundation model on single-cell transcriptomics. Nature Methods 21, 1481–1491 (2024).

[26] Cui, H. et al. scGPT: toward building a foundation model for single-cell multi-omics using generative AI. Nature Methods 21, 1470–1480 (2024).

[27] Theodoris, C. V. et al. Transfer learning enables predictions in network biology. Nature 618, 616–624 (2023).

[28] Rosen, Y. et al. Universal Cell Embeddings: A foundation model for cell biology. bioRxiv2023–11 (2023).

[29] Adduri, A. K. et al. Predicting cellular responses to perturbation across diverse contexts with State. bioRxiv 2025–06 (2025).

[30] Li, C., Wei, L. & Zhang, X. Unified multimodal learning enables generalized cellular response prediction to diverse perturbations. bioRxiv 2025–11 (2025).

[31] Yuan, X. et al. PerturbDiff: Functional diffusion for single-cell perturbation modeling. arXiv preprint arXiv:2602.19685 (2026).

[32] Chuai, G. et al. Towards building a world model to simulate perturbation-induced cellular dynamics by AlphaCell. bioRxiv 2026–03 (2026).

[33] Wang, C. et al. X-Cell: Scaling causal perturbation prediction across diverse cellular contexts via diffusion language models. bioRxiv 2026–03 (2026).

[34] Zhu, O. & Li, J. Scouter predicts transcriptional responses to genetic perturbations with large language model embeddings. Nature Computational Science 1–8 (2025).

[35] Chen, Y. & Zou, J. GenePert: Leveraging GenePT embeddings for gene perturbation prediction. bioRxiv 2024–10 (2024).

[36] Jiang, Q. et al. Phenotype-guided in silico molecular generation using large language models. bioRxiv 2026–01 (2026).

[37] Wu, Y. et al. PerturBench: Benchmarking machine learning models for cellular perturbation analysis. Advances in Neural Information Processing Systems 38 (2026).

[38] Wei, Z. et al. Benchmarking algorithms for generalizable single-cell perturbation response prediction. Nature Methods 23, 451–464 (2026).

[39] Evan, G. I. & Vousden, K. H. Proliferation, cell cycle and apoptosis in cancer. Nature 411, 342–348 (2001).

[40] McFarland, J. M. et al. Multiplexed single-cell transcriptional response profiling to define cancer vulnerabilities and therapeutic mechanism of action. Nature Communications 11, 4296 (2020).

[41] Zhang, Z., Li, T. & Zhou, P. Learning stochastic dynamics from snapshots through regularized unbalanced optimal transport. The Thirteenth International Conference on Learning Representations (2025). URL https://openreview.net/forum?id=gQlxd3Mtru.

[42] Peng, Q. et al. WFR-FM: Simulation-free dynamic unbalanced optimal transport. The Fourteenth International Conference on Learning Representations (2026).

[43] Zdrazil, B. et al. The ChEMBL database in 2023: a drug discovery platform spanning multiple bioactivity data types and time periods. Nucleic Acids Research 52, D1180–D1192 (2024).

[44] Souers, A. J. et al. ABT-199, a potent and selective BCL-2 inhibitor, achieves antitumor activity while sparing platelets. Nature Medicine 19, 202–208 (2013).

[45] Carter, B. Z. et al. Combined targeting of BCL-2 and BCR-ABL tyrosine kinase eradicates chronic myeloid leukemia stem cells. Science Translational Medicine 8, 355ra117 (2016).

[46] Jordan, M. A. & Wilson, L. Microtubules as a target for anticancer drugs. Nature Reviews Cancer 4, 253–265 (2004).

[47] Andre, F. et al. Alpelisib for PIK3CA-mutated, hormone receptor-positive advanced breast cancer. New England Journal of Medicine 380, 1929–1940 (2019).

[48] Lo-Coco, F. et al. Retinoic acid and arsenic trioxide for acute promyelocytic leukemia. New England Journal of Medicine 369, 111–121 (2013).

[49] Tse, C. et al. ABT-263: a potent and orally bioavailable Bcl-2 family inhibitor. Cancer Research 68, 3421–3428 (2008).

[50] Shi, J., Zhou, Y.Huang, H.-C. & Mitchison, T. J. Navitoclax (ABT-263) accelerates apoptosis during drug-induced mitotic arrest by antagonizing bcl-xl. Cancer Research 71, 4518–4526 (2011).

[51] Goff, D. J. et al. A pan-bcl2 inhibitor renders bone-marrow-resident human leukemia stem cells sensitive to tyrosine kinase inhibition. Cell stem cell 12, 316–328 (2013).

[52] Oesinghaus, L. et al. A single-cell cytokine dictionary of human peripheral blood. bioRxiv(2025).

[53] Sha, Y., Qiu, Y., Zhou, P. & Nie, Q. Reconstructing growth and dynamic trajectories from single-cell transcriptomics data. Nature Machine Intelligence 6, 25–39 (2024).

[54] Peng, Q., Zhou, P. & Li, T. stVCR: spatiotemporal dynamics of single cells. Nature Methods23, 542–553 (2026).

[55] Wang, D. et al. Joint velocity-growth flow matching for single-cell dynamics modeling. Advances in Neural Information Processing Systems 38, 160552–160588 (2026).

[56] Wang, X., Wang, R., Peng, Q., Zhou, P. & Li, T. WFR-MFM: One-step inference for dynamic unbalanced optimal transport. arXiv preprint arXiv:2601.20606 (2026).

[57] Wagner, D. E. & Klein, A. M. Lineage tracing meets single-cell omics: opportunities and challenges. Nature Reviews Genetics 21, 410–427 (2020).

[58] Lukinavičius, G. et al. Fluorogenic probes for live-cell imaging of the cytoskeleton. Nature Methods 11, 731–733 (2014).

[59] Dhainaut, M. et al. Spatial CRISPR genomics identifies regulators of the tumor microenvironment. Cell 185, 1223–1239 (2022).

[60] Binan, L. et al. Simultaneous CRISPR screening and spatial transcriptomics reveal intracellular, intercellular, and functional transcriptional circuits. Cell 188, 2141–2158 (2025).

[61] Shen, K. et al. Spatial perturb-seq: single-cell functional genomics within intact tissue architecture. Nature Communications 17, 3018 (2026).

[62] Zhang, H. et al. Uncovering spatially resolved functional genomics with CRISPR screen sequencing. Cell (2026).

[63] Baysoy, A. et al. Large-scale, spatially resolved panoramic CRISPR screening in native tissue environments using Perturb-DBiT. Nature Biotechnology (2026).

[64] Jin, S. et al. Inference and analysis of cell-cell communication using CellChat. Nature Communications 12, 1088 (2021).

[65] Liero, M., Mielke, A. & Savaré, G. Optimal transport in competition with reaction: The Hellinger–Kantorovich distance and geodesic curves. SIAM Journal on Mathematical Analysis 48, 2869–2911 (2016).

[66] Chizat, L., Peyré, G., Schmitzer, B. & Vialard, F.-X. An interpolating distance between optimal transport and Fisher–Rao metrics. Foundations of Computational Mathematics 18, 1–44 (2018).

[67] Kondratyev, S., Monsaingeon, L. & Vorotnikov, D. A new optimal transport distance on the space of finite Radon measures. Advances in Differential Equations 21, 1117–1164 (2016).

[68] Geng, Z., Deng, M., Bai, X., Kolter, Z. & He, K. Mean flows for one-step generative modeling. Advances in Neural Information Processing Systems 38, 75460–75482 (2026).

[69] Perez, E., Strub, F., De Vries, H., Dumoulin, V. & Courville, A. FiLM: Visual reasoning with a general conditioning layer. Proceedings of the AAAI conference on artificial intelligence 32 (2018).

[70] Peebles, W. & Xie, S. Scalable diffusion models with transformers. Proceedings of the IEEE/CVF International Conference on Computer Vision 4195–4205 (2023).

[71] Flamary, R. et al. POT: Python optimal transport. Journal of Machine Learning Research22, 1–8 (2021).

[72] Virshup, I. et al. The scverse project provides a computational ecosystem for single-cell omics data analysis. Nature Biotechnology 41, 604–606 (2023).

[73] Virshup, I., Rybakov, S., Theis, F. J., Angerer, P. & Wolf, F. A. anndata: Access and store annotated data matrices. Journal of Open Source Software 9, 4371 (2024).

[74] Wolf, F. A., Angerer, P. & Theis, F. J. SCANPY: large-scale single-cell gene expression data analysis. Genome Biology 19, 15 (2018).

[75] Qiu, X. et al. Mapping transcriptomic vector fields of single cells. Cell 185, 690–711 (2022).

[76] Ashburner, M. et al. Gene Ontology: tool for the unification of biology. Nature Genetics 25, 25–29 (2000).

[77] Subramanian, A. et al. Gene set enrichment analysis: A knowledge-based approach for interpreting genome-wide expression profiles. Proceedings of the National Academy of Sciences 102, 15545–15550 (2005).

